# Benchmarking macaque brain gene expression for horizontal and vertical translation

**DOI:** 10.1101/2024.08.18.608440

**Authors:** Andrea I. Luppi, Zhen-Qi Liu, Justine Y. Hansen, Rodrigo Cofre, Elena Kuzmin, Seán Froudist-Walsh, Nicola Palomero-Gallagher, Bratislav Misic

## Abstract

The spatial patterning of gene expression shapes cortical organization and emergent function. Advances in spatial transcriptomics make it possible to comprehensively map cortical gene expression in both humans and model organisms. The macaque is a particularly valuable model organism, due to its evolutionary similarity with the human. The translational potential of macaque gene expression rests on the assumption that it is a good proxy for spatial patterns of corresponding proteins (vertical translation) and for spatial patterns of ortholog human genes (horizontal translation). Here we systematically benchmark the spatial distribution of gene expression in the macaque cortex against (a) cortical receptor density in the macaque and (b) cortical gene expression in the human. We find that there is moderate cortex-wide correspondence between gene expression and protein density in the macaque, which is improved by considering layer-specific gene expression. We find greater correspondence between orthologous gene expression in the macaque and human. Inter-species correspondence of gene expression exhibits systematic regional heterogeneity, with greater correspondence in unimodal than transmodal cortex, mapping onto patterns of evolutionary cortical expansion. We extend these results to additional micro-architectural features using macaque immunohistochemistry and T1w:T2w ratio, and replicate them using macaque RNA-seq and human RNA-seq gene expression. Collectively, the present results showcase both the potential and limitations of macaque spatial transcriptomics as an engine of translational discovery within and across species.

## INTRODUCTION

The macaque is a foundational and widely used model organism in neuroscience [5, 19, 55, 79, 80, 85, 95]. Animal models allow invasive tracing, imaging and recording, as well as numerous experimental manipulations that would not be possible in humans, allowing scientists to address a wider range of scientific questions [15, 84]. Compared to rodents and other non-human primates (NHPs) such as marmosets, the macaque possesses several desirable features for studying human brain structure and function. These include shared evolutionary history, gyrified cortex, and a more humanlike behavioural repertoire including greater ability to exert cognitive control against distractors during cognitive performance [62, 85, 89, 95]. These similarities are grounded in the genetic relatedness between the two species (92% genetic alignment with *Homo sapiens*) [34]. As a result, the macaque has been used to study the evolutionary and developmental origins of brain anatomy, cognition, and behaviour [88, 114], as well as the consequences of targeted genetic mutations in transgenic animals [36, 61, 70, 129].

Recent advances in high-throughput sequencing make it possible to map spatial patterns of gene expression across the cortex of humans and other species. This is valuable because gene expression is heterogeneous across the cortex: spatial patterns of gene expression provide a blueprint for the brain’s structural and functional architecture [12, 22, 54, 67]. Indeed, multiple reports have inferred transcriptional signatures of cell types [12, 101, 103, 109, 121], neurotransmitter receptors [4, 13, 17, 29, 90, 110, 132], laminar differentiation [122], cortical thickness [12, 127], structural and functional connectivity [72, 92, 96, 117], brain dynamics [30, 106], cognitive specialization [51] development [116, 127] and disease [52, 101, 133], among others [87]. Key to this endeavour has been the development of comprehensive spatial transcriptomics datasets. However, until recently databases of comprehensive cortical gene expression have been restricted to the human [54] and mouse [21, 67]. An exciting recent development is the availability of regionally-resolved transcriptomics for the macaque brain [12, 22, 23], providing an unprecedented opportunity to combine this species’ experimental accessibility and genetic similarity to humans [78].

Effective translational discovery from macaque gene expression depends on two fundamental questions. The first question is whether orthologous genes display similar spatial patterning across human and macaque cortex. In other words, can regionally specific gene expression findings in the macaque be extrapolated to the human? The second question is whether spatial patterns of protein-coding genes in the macaque are a good proxy for spatial patterns of protein density in the same species, given the numerous intervening steps between mRNA transcription and expression of the corresponding protein at its final location. In other words, if we know the regional expression pattern of a gene from the macaque brain, can we infer the regional distribution of the protein that this gene codes for? Ultimately, as the field embarks towards comprehensive transcriptional mapping of the macaque brain, it is necessary to benchmark both the horizontal translational potential of these datasets (from macaque gene expression to human gene expression) [78] as well as their vertical translational potential (from macaque gene expression to other modalities within the macaque brain) [78].

Here, we address these questions by analysing a recently released database of *Macaca fascicularis* cortical gene expression [22] from high-resolution, large-fieldof view spatial transcriptomics (“stereo-seq”) [21]. To benchmark horizontal translation from macaque to human, we compare macaque stereo-seq transcriptomics with human cortical gene expression from post-mortem microarray data of six adult donors’ brains, made available by the Allen Human Brain Atlas (AHBA) [54, 75]. To benchmark vertical translation between different modalities, we compare macaque cortical gene expression [22] against measurements of receptor density in the cortex of *Macaca fascicularis* obtained from post-mortem autoradiography [39] (Fig. 1). Specifically, we focus on neurotransmitter receptors, a class of protein complexes that are essential for brain function, and therefore of particular interest in both basic research and clinical applications. Assessing the correspondence between gene expression and receptor density is of particular interest because gene expression is often used as a proxy for receptor density in the brain [4, 13, 16, 17, 20, 29, 41, 43, 58, 90, 110, 132]. However, measurements of gene expression and protein density do not always align [8, 50, 63, 64, 81, 82, 131, 136]. In the present report we address these fundamental questions by assessing the extent to which gene expression in the macaque cortex reflects human gene expression, and the extent to which it reflects protein availability in macaque.

**Figure 1.**
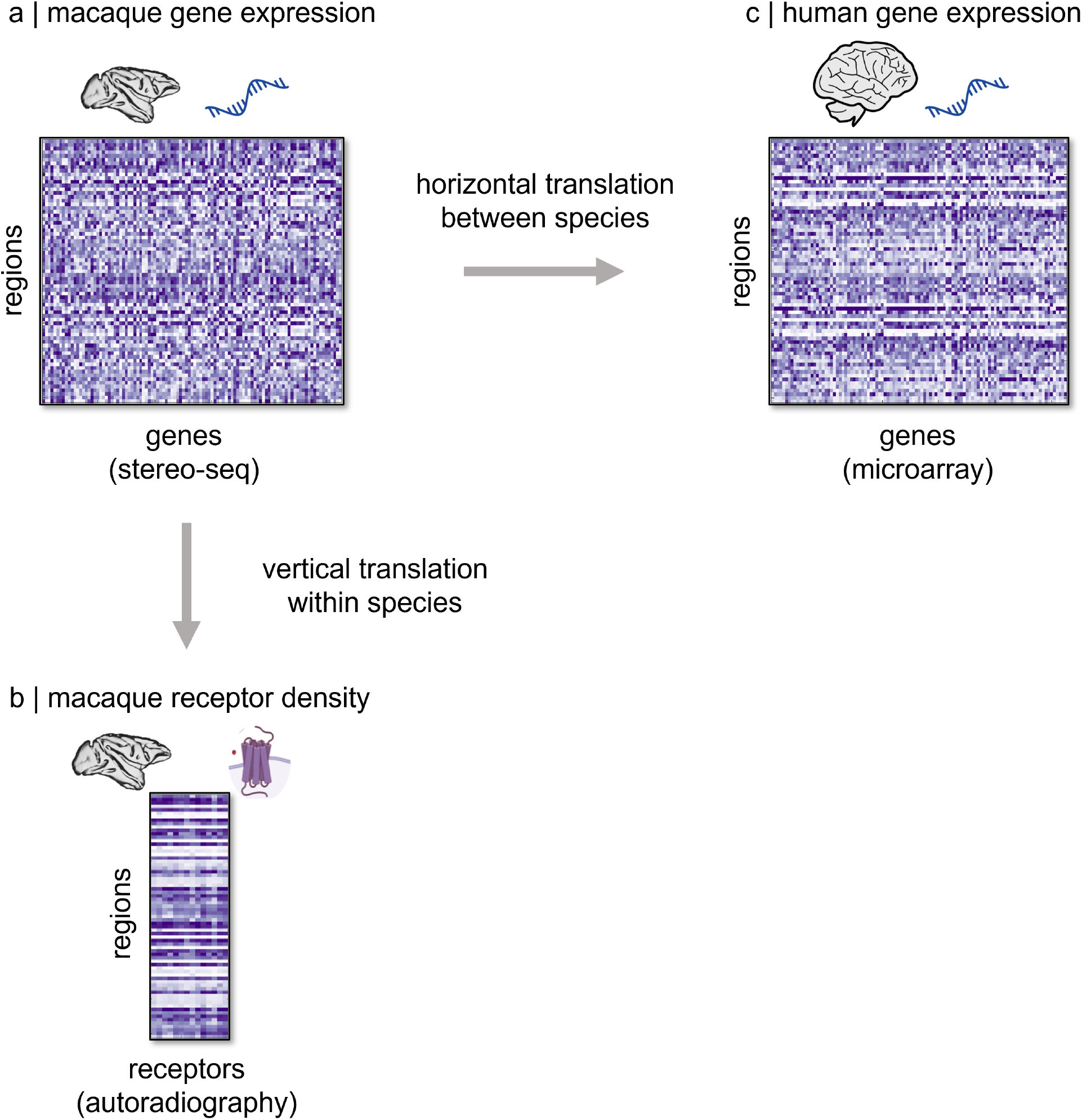
Overview. We compare macaque cortical gene expression from stereo-seq [22] with human gene expression from microarray in homologous regions [54], and with macaque density of receptors per neuron quantified from autoradiography [39]

## RESULTS

We analyze three openly-available datasets:

- Macaque cortical “stereo-seq” gene expression [22].
- Macaque cortical receptor density per neuron from *in vitro* autoradiography [39].
- Human cortical microarray gene expression [54].

To obtain a common space for comparison, we apply the canonical Regional Mapping (RM) parcellation of the macaque cortex developed by Kötter and Wanke [9, 66], alongside its recent translation to the human brain [10], allowing us to obtain a one-to-one mapping between regions in the two species (Fig. S1; see *Methods* for details). We then ask two questions. First: what is the spatial correspondence between macaque gene expression and neurotransmitter receptor density? Second: what is the spatial correspondence between macaque cortical gene expression, and human cortical gene expression?

### Gene-receptor correspondence in the macaque

To determine whether gene expression is a suitable proxy for receptor density per neuron across macaque cortex, we compare stereo-seq macaque gene expression with quantitative measurements of corresponding receptor density from *in vitro* autoradiography, which uses radioactive ligands to quantify the endogenous receptors bound in the cell membrane [22, 39]. We focus on thirteen receptors, covering both ionotropic and metabotropic receptors and spanning 6 neurotransmitter systems: noradrenergic (*α*_*1*_, *α*_*2*_), serotonergic (*5HT*_*1A*_, *5HT*_*2A*_), dopaminergic (*D*_*1*_), cholinergic (*M*_*1*_, *M*_*2*_), glutamatergic (*AMPA, kainate, NMDA*) and GABAergic (*GABA*_*A*_, *GABA*_*B*_, *GABA*_*A/BZ*_). For each receptor, we identify the corresponding genes in the stereo-seq expression data. For details on how genes were selected for multimeric receptors, see *Methods*; the complete set of genes related to receptor subunits is shown in Fig. S2. Fig. 2 shows the spatial correlation between receptor density per neuron (x-axis) and gene expression (y-axis) for each receptor. All correlations are evaluated against spatial autocorrelation-preserving nulls (see *Methods*), and significant associations (*p <* 0.05) are shown in indigo. We find that few macaque neurotransmitter receptors correlate significantly with the expression of their main corresponding genes (5 out of 18 or 28%; Fig. 2) A similar result is observed when considering the full set of receptor subunits (16 out of 55, or 29.1%; Fig. S2. The observation of highly variable gene-receptor correspondence is consistent with findings in humans, using both *in vivo* positron emission tomography (PET) and postmortem autoradiography [8, 50, 63, 64, 81, 82, 136].

**Figure 2.**
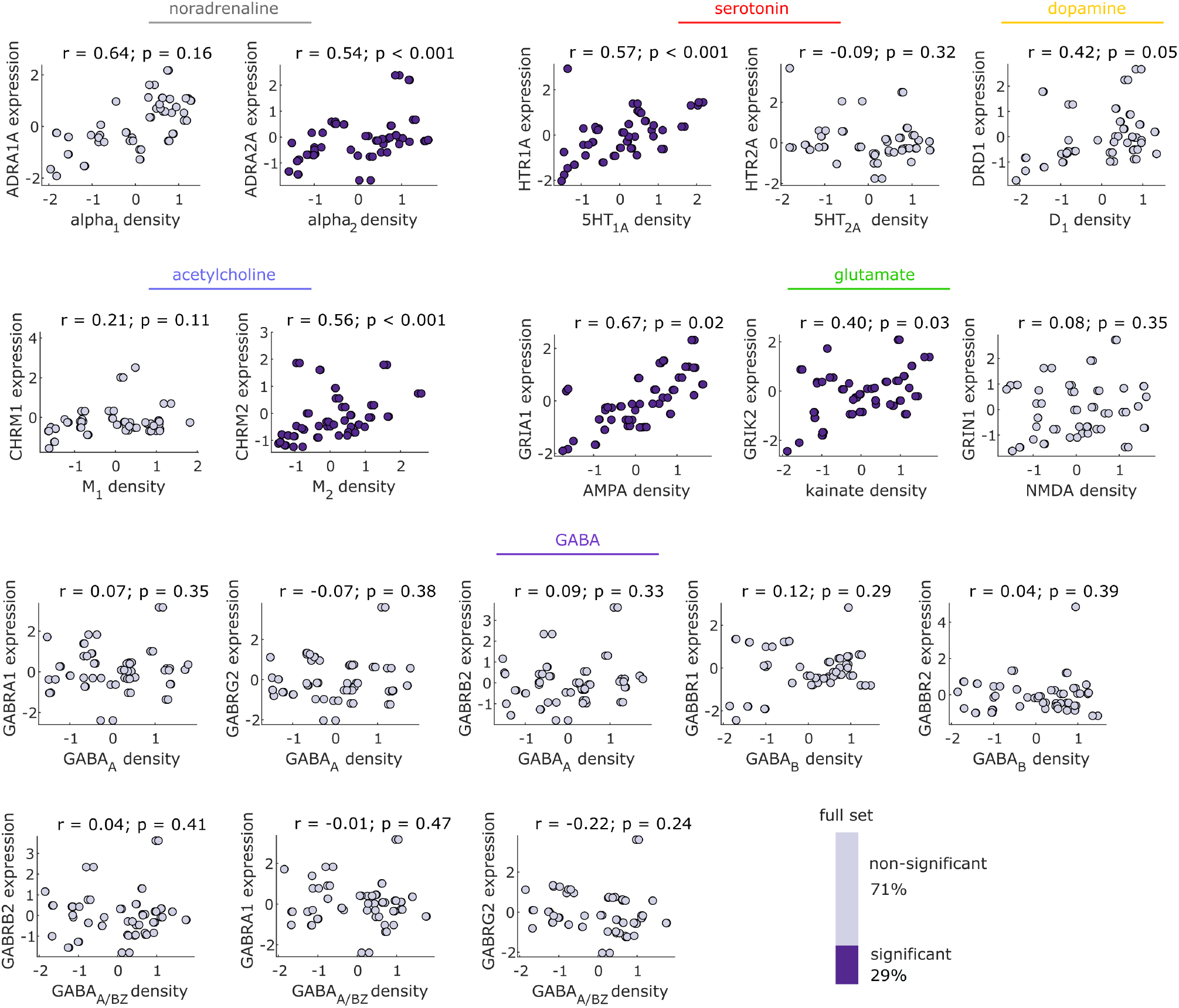
Autoradiography-derived receptor densities versus gene expression in the macaque. The density of only few neurotransmitter receptors correlates significantly with the expression of their corresponding gene, across macaque cortex. Indigo scatter plots indicate significant positive gene-receptor correspondence (Spearman’s *r, p <* 0.05 against a null distribution with preserved spatial autocorrelation). Values are z-scored. Bars indicates the proportion of significant correlations (indigo) out of all gene-receptor pairs (grey), for the subset shown, and for the full set in Fig. S2.

For receptors pertaining to neuromodulatory systems, we observe significant correlations between density per neuron of the adrenergic *α*_*2*_ receptor with *ADRA2A* gene expression; serotonergic *5HT*_*1A*_ receptor density and *HTR1A* gene expression; and muscarinic acetylcholine receptor *M*_*2*_ and *CHRM2* gene expression. Notably, *DRD1* gene expression barely fails to meet the threshold for a statistically significant correlation with dopamine *D*_*1*_ receptor density, after controlling for spatial autocorrelation (*p* = 0.05); similarly, *α*_*1*_ receptor density and *ADRA1A* gene expression exhibit the second-highest value of correlation (Spearman’s *r* = 0.67), albeit not significant after accounting for spatial autocorrelation. For glutamate receptors, we observe significant correlations between *AMPA* receptor density and *GRIA1, GRIA3* and *GRIA4* gene expression; kainate receptor density and *GRIK1, GRIK2* and *GRIK3* gene expression; *NMDA* receptor density and *GRIN2C* and *GRIN3A* gene expression. For GABA receptors, we observed significant correlations between *GABA*_*A*_ receptor density and expression of *GABRA6* and *GABRQ* genes; (with *p* = 0.05 for *GABRA5*); and between *GABA*_*A/BZ*_ binding site density and expression of *GABRA5* and *GABRQ* (Fig. S2).

Notably, the correspondence between *5HT*_*1A*_ receptor density and *HTR1A* gene expression had also been observed in humans with both *in vivo* PET and *post-mortem* autoradiography, by multiple studies [8, 50, 63, 81], and even between human genes and macaque receptors [39]. Altogether, the overall level of correspondence between gene expression and receptor density is consistent with recent findings in the human brain [50, 63, 64, 81, 136].

One potential reason why receptor density may not align perfectly with gene expression is that gene transcription occurs in the soma, whereas many receptors are expressed at the synapse, which for presynaptic receptors may be far away for neurons with long axons. To investigate this possibility, we correlate the receptor density of each cortical region against the average of the gene expression of regions that are directly connected to it by efferent white matter tracts, weighted by the strength of the connection (see *Methods*) [52, 104, 105]. The rationale for this approach is that if receptors expressed in one region are the result of genes transcribed at the other end of long-range axons in other regions, then the receptor’s density should be better predicted by considering the average of gene expression in neighbouring regions (weighted by the weight of the connection). Our results rule out this possibility (Fig. S3). No new significant gene-receptor correlations emerge, and the only significant correlations are between *HTR1A* gene expression and *5HT*_*1A*_ receptor density, and between *GABRA6* gene expression and *GABA*_*A*_ receptor density (Fig. S3), which were also significant in the original analysis.

Since different neuron types can be preferentially localised in specific cortical layers, we next seek to determine whether the imperfect gene-receptor correlations could be attributable to layer-specificity of relationships between gene transcription, and the expression of the corresponding proteins. Therefore, we repeat our previous analysis, but using layer-specific gene expression data (Fig. S4 – S9). This approach also allows us to evaluate differences between layers. We find that the number of gene-receptor pairs exhibiting significant spatial correlation across at least one layer is 33 out of 55 (60%): significantly higher than the 28% (16 gene-receptor pairs) that exhibit significance when aggregating across layers (Fig. 3a-c; *χ*^2^ = 10.64; *p* = 0.001). Of these 33 significant gene-receptor pairs, 28 (51%) have significant correlations for 2 or more layers. Crucially, the distribution is not uniform across layers, with layers 4 and 6 enjoying relatively higher proportion of significant correlations for the glutamate receptors, and layers 3 and 5 for the receptors pertaining to neuromodulatory systems (Fig. 3d). Notably, differences also emerge among receptor types: only 18.7% of all layer-wise gene-receptor pairs for GABA are significantly correlated, which is a significantly lower proportion than observed for both glutamate receptors (39%; *χ*^2^ = 13.19; *p* = 0.003), and receptors pertaining to neuromodulatory systems (33%; *χ*^2^ = 4.81; *p* = 0.028). This layer-specific pattern of correspondences between genes and receptors showcases the intertwined nature of cortical chemoarchitecture and laminar differentiation, highlighting the need for more comprehensive datasets of layer-specific receptor density.

**Figure 3.**
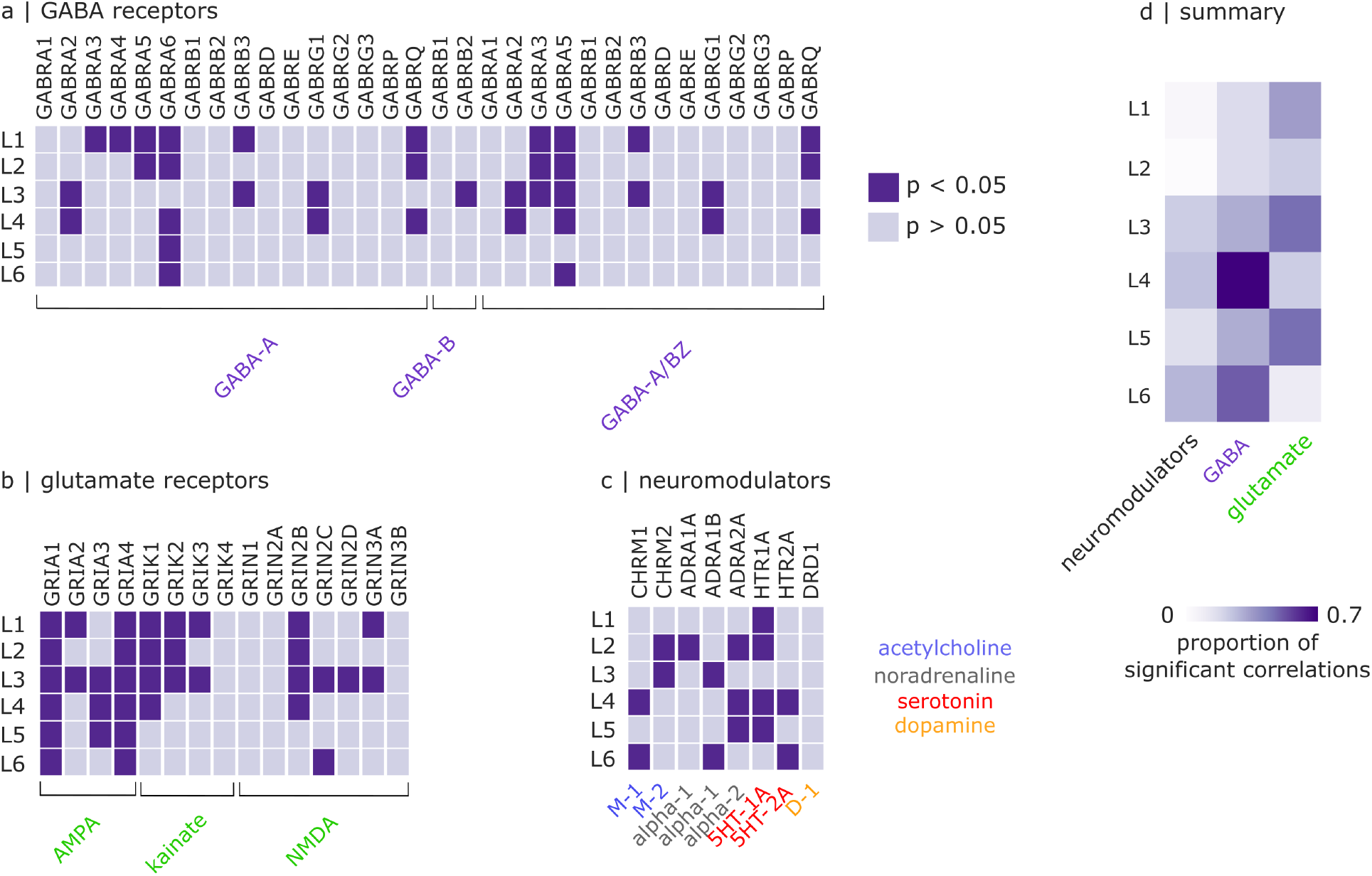
Significance between macaque gene expression, and macaque receptor expression across cortical layers and receptor types. (**a**) Significance of gene-receptor correlations for *GABA* receptors. (**b**) Significance of gene-receptor correlations for glutamate receptors. (**c**) Significance of gene-receptor correlations for receptors pertaining to neuromodulatory systems (acetylcholine, noradrenaline, serotonin, dopamine). For a-c, indigo cells indicate significant positive gene-receptor correspondence (Spearman’s *r, p <* 0.05 against a null distribution with preserved spatial autocorrelation); grey cells indicate no significance. Columns indicate cortical layers, and rows indicate gene-receptor pairs.(**d**) Summary of the proportion of significant correlations from a-c, for each layer and each broad receptor type.

### Macaque gene expression recapitulates micro-architectural features

Is moderate gene-protein correspondence due to noisy gene expression data? Does macaque gene expression align with other micro-architectural features? One way to evaluate this possibility is by assessing whether gene-protein correlations that we should expect to observe, based on the literature, can be recovered using these data. This is indeed the case for the correspondence between *5HT*_*1A*_ receptor density and *HTR1A* gene, which has been reported in humans with both *in vivo* PET and *post-mortem* autoradiography [50, 63, 81]. Notably, Fulcher et al. [41] reported high within-species correspondence in the mouse between *Pvalb* gene expression and the density of neurons expressing parvalbumin (the protein that this gene codes for), as well as high correspondence between mouse *Pvalb* and *Calb2* and human *PVALB* and *CALB2* gene expression. In addition, Burt et al. [16] observed that macaque protein density of the calcium-binding proteins parvalbumin and calretinin from immunohistochemistry exhibit similar spatial patterns as the expression of human *PVALB* and *CALB2* genes, which code for these proteins. Taken together, these previous results suggest that we should expect to observe similar spatial patterns for macaque *PVALB* gene expression and parvalbumin density, and for macaque *CALB2* expression and calretinin protein density.

Indeed, we show that macaque *PVALB* gene expression is significantly correlated (*r* = 0.37, *p <* 0.001) with an independent measure of density of parvalbumin-immunoreactive interneurons from immunohistochemistry [26, 31, 42, 65] assembled and made available by Burt et al. [16] (Fig. 4a). Likewise, macaque *CALB2* gene expression is significantly spatially correlated with density of calretinin-immunoreactive interneurons in the macaque cortex [16] (*r* = 0.65, *p <* 0.001; Fig. 4a). We replicate this result using the relative prevalence of calretinin-immunoreactive and parvalbumin-immunoreactive neurons across a subset of visual, auditory, and somatosensory regions of the macaque from Kondo et al. [65], one of the original studies aggregated by Burt and colleagues (Fig. S10).

**Figure 4.**
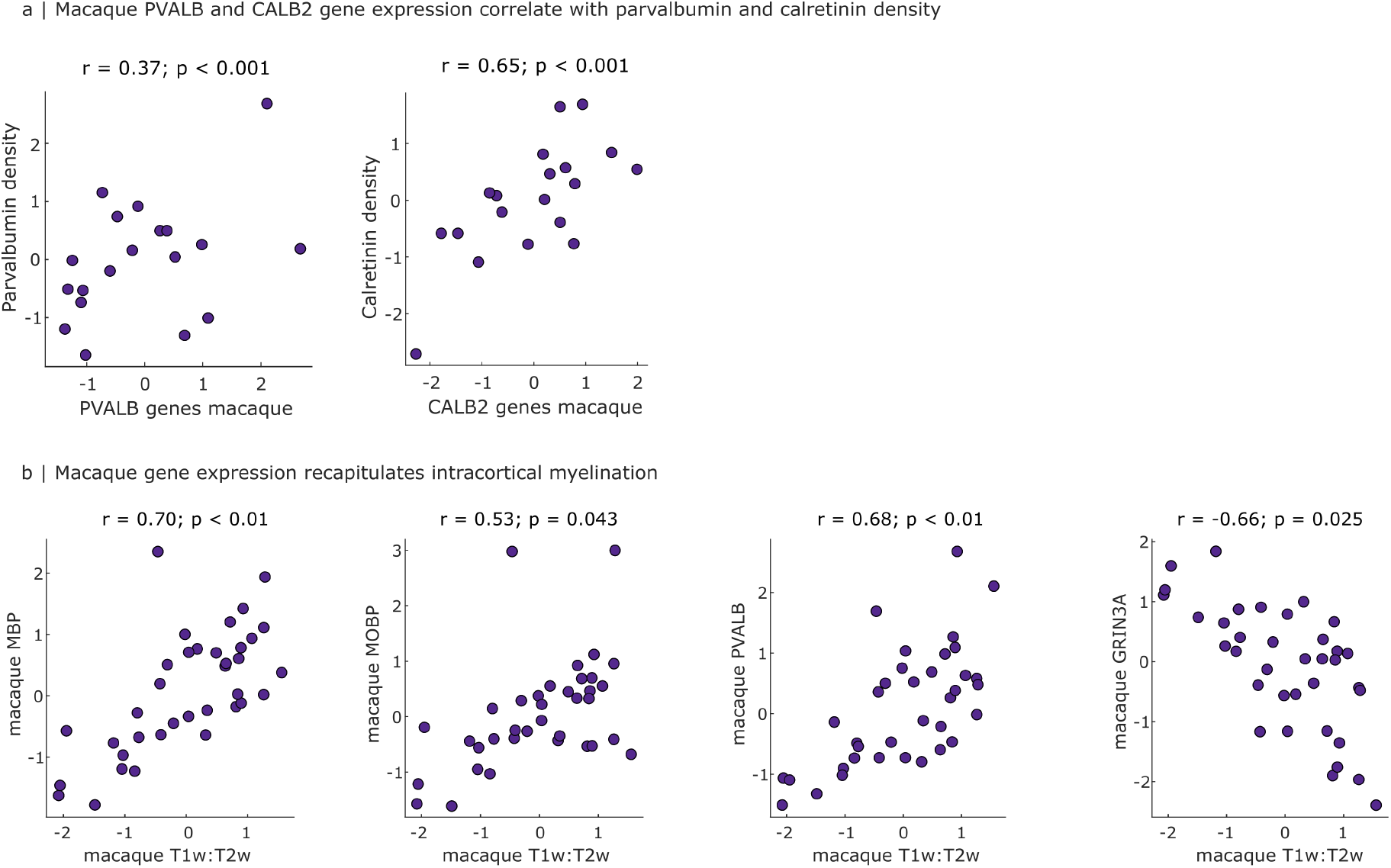
Macaque gene expression recapitulates micro-architectural features. (**a**) Spatial correspondence of macaque *PVALB* and *CALB2* gene expression, with parvalbumin and calretinin protein density. (**b**) Spatial correspondence of macaque gene expression with intracortical myelination from T1w:T2w ratio. Indigo scatter plots indicate significant spatial correspondence (Spearman’s *r, p <* 0.05 against a null distribution with preserved spatial autocorrelation). Values are z-scored.

Additionally, Fulcher et al. [41] reported that both humans and mice exhibit significant spatial correspondence between *in vivo* intracortical myelination (T1w:T2w ratio map), and expression of the key myelin-related genes *MBP/Mbp* and *MOBP/Mobp*, as well as *PVALB/Pvalb* and *GRIN3A/Grin3a*. Here, we demonstrate that each of these relationships is also observed in the macaque: we find significant positive spatial correlations between macaque intracortical myelination, and macaque cortical expression of *MBP* (*r* = 0.70, *p <* 0.001) *MOBP* (*r* = 0.53, *p* = 0.043), and *PVALB* (*r* = 0.68, *p <* 0.001); as well as a significant negative association between T1w:T2w ratio and *GRIN3A* gene expression (*r* = *−*0.66, *p* = 0.025) (Fig. 4b) – precisely as reported in [40]. In addition to demonstrating a close spatial correspondence between myelin-related gene expression and an *in vivo* marker of cortical myelination in the macaque [45, 46, 59], these results also demonstrate close alignment of our results with two different mammalian species. Collectively, despite moderate spatial alignment in the specific case of gene-receptor correspondence, these results indicate that macaque gene expression can recapitulate many other *in vivo* and *ex vivo* features of macaque cortical microarchitecture.

### Cross-species correspondence of gene expression

To assess the translational potential of macaque gene expression to human, we next compare macaque spatial patterns of gene expression, against patterns of human gene expression obtained from the Allen Institute’s microarray data [54]. We focus on genes related to receptor subunits, as well as cell-type markers (parvalbumin, somatostatin, calbindin, vasoactive intestinal polypeptide) and the four most abundant mRNAs in myelin (*MBP, FTH1, PLEKHB1*, and *MOBP*) by combining the lists of receptor-related genes from [50, 81] with [41], as well as genes coding for SLC6 transporters, histidine decarboxylase (*HDC*), and syntaxin (*STX1A*), hyperpolarization-activated cyclic nucleotide-gated channels (HCN), potassium channels (KCN), and sodium voltage-gated channels (SCN), which are also relevant for brain function. For all analyses, we only consider genes that (a) have a macaque ortholog, and (b) are available in both the human and macaque datasets after accounting for all preprocessing criteria, such as intensity filtering, yielding a total of 106 genes.

We find that 52% of genes considered (55*/*106) exhibit significant spatial correlation between humans and macaques, after accounting for spatial autocorrelation. A representative sample of cross-species gene correlations pertaining to receptors is shown in Fig. 5; the full set is shown in Fig. S11. In other words, we find that interspecies correspondence of gene expression for genes pertaining to cell types and receptor subunits is greater than the correspondence between macaque gene expression and density of the corresponding receptors, as shown in the previous section (*χ*^2^ = 7.63, *p <* 0.006). However, when we consider macaque gene-receptor pairs having at least one significant correlation across cortical layers (instead of aggregating over layers), then the proportion (60%) is comparable to the proportion of significant inter-species correlations of gene expression (51%) (*χ*^2^ = 0.96, *p* = 0.327).

**Figure 5.**
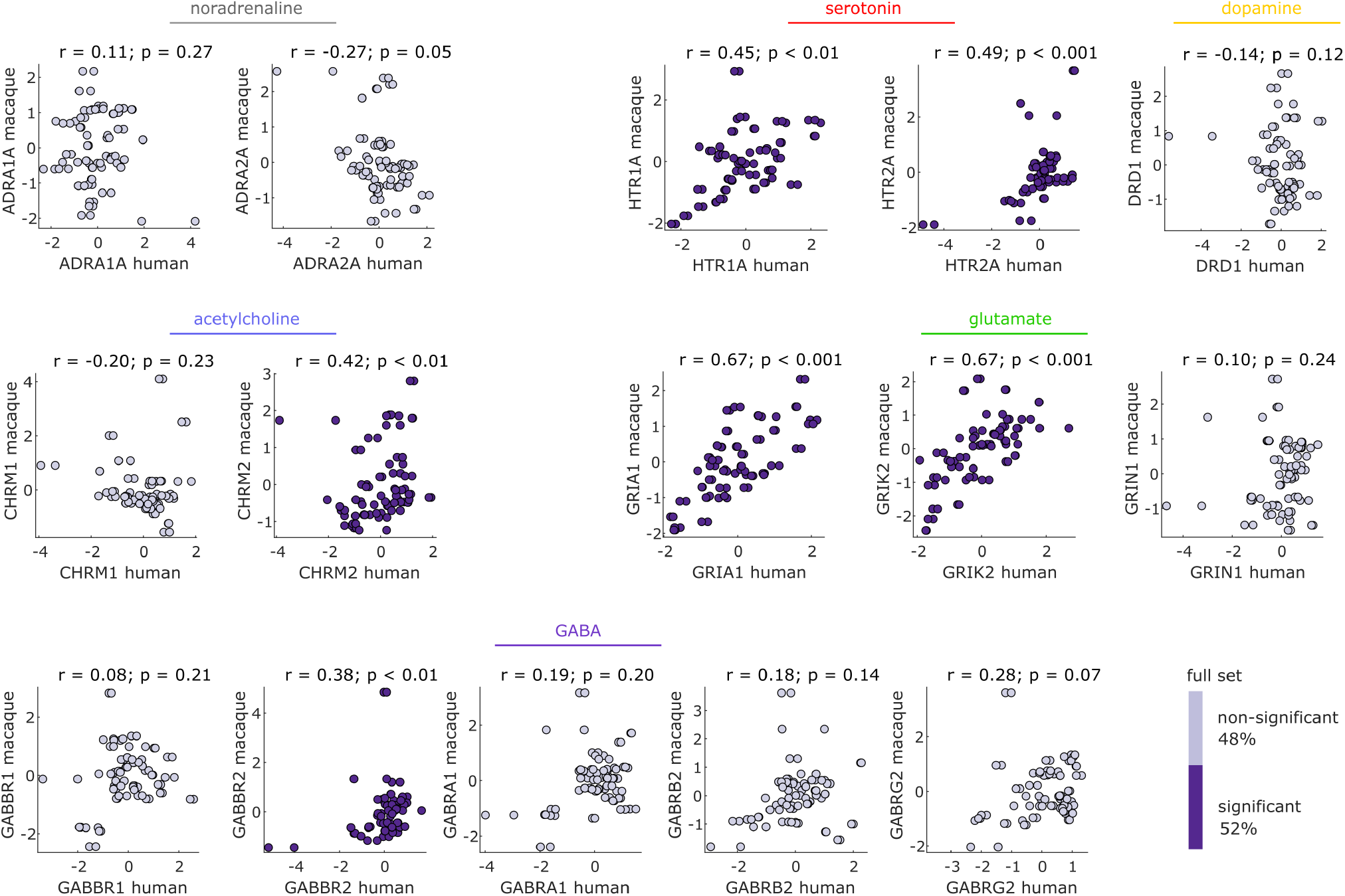
Spatial correspondence between the expression of the main available brain-relevant genes in humans (from microarray) and macaques (from stereo-seq) The majority of macaque genes exhibit significant spatial correlation with the corresponding human ortholog genes. Indigo scatter plots indicate significant human-macaque correspondence (Spearman’s *r, p <* 0.05 against a null distribution with preserved spatial autocorrelation). Values are z-scored. Bar indicates the proportion of significant correlations (indigo) out of all gene-gene pairs (grey).

Is inter-species gene expression correspondence uniform across the brain or is it regionally heterogeneous? Complementary to correlating expression of each gene across regions (i.e., each data-point being a cortical region), we can also consider the correlation of expression of different genes at each cortical region (i.e., with data-points being genes) (Fig. 6a). We find that the regional inter-species correlation of genes is significantly greater in unimodal compared to transmodal regions of the cortex (Fig. 6a). These systematic differences between gene expression in the two species are reflected in regionally heterogeneous phylogenetic divergence between humans and nonhuman primates, including macaques [32, 35, 56, 71, 124, 125, 128]. Indeed, cortical regions that exhibit the lowest correspondence of gene expression between macaque and human, are also the regions that have undergone the greatest cortical expansion between the two species, as quantified in a recent report of human-macaque cortical expansion by Xu and colleagues [128]. Specifically, there is a significant negative spatial correlation between the two cortical patterns: *Spearman*^*′*^*sr* = *−*0.35, *p* = 0.002 (Fig. 6b).

**Figure 6.**
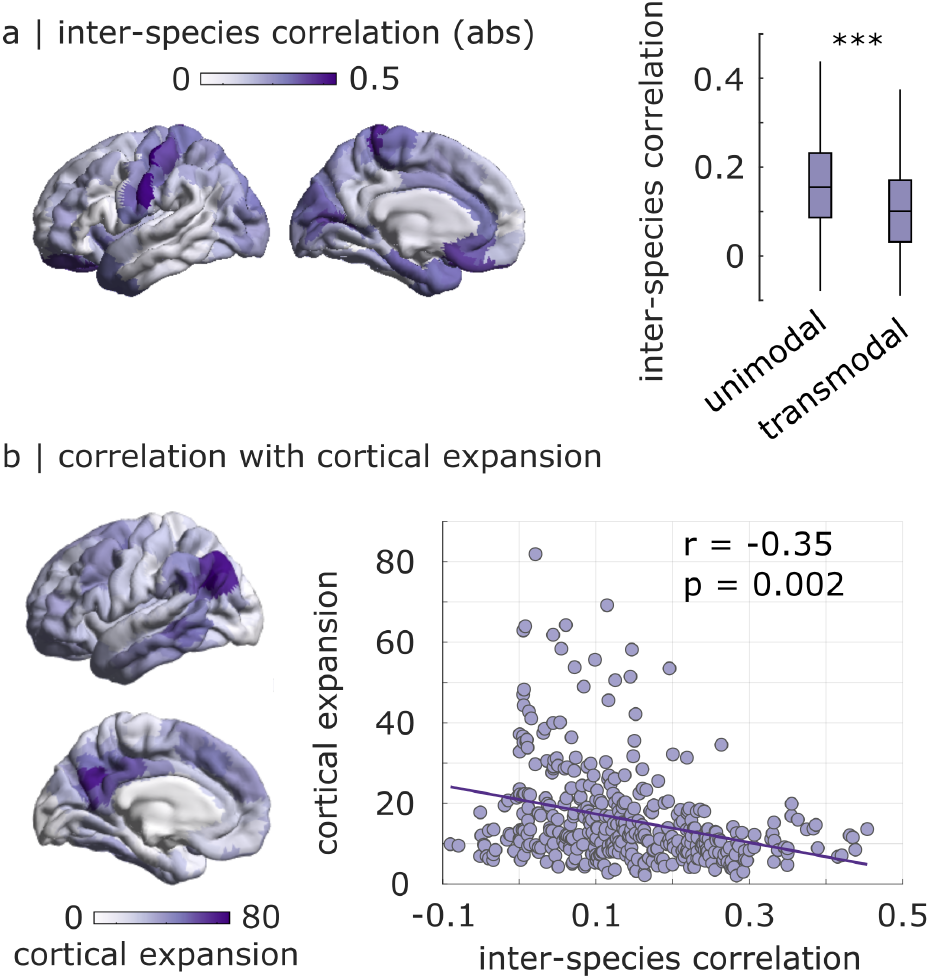
Regional correlation between human and macaque patterns of cortical gene expression recapitulates cortical expansion. (**a**) Intensity of color of each region indicates the magnitude of Spearman correlation between human microarray and macaque stereo-seq gene expression for that region, across genes. Data are re-parcellated from the human version of the Regional Mapping atlas to the Schaefer-400 parcellation. The inter-species correlation of regional gene expression is significantly higher in unimodal than transmodal regions; unimodal mean = 0.16; transmodal mean = 0.11; *t*(398) = 5.20, Cohen’s *d* = 0.52, *p <* 0.001 from independent-samples t-test. (**b**) The cortical pattern of regional inter-species correlations of gene expression is significantly negatively correlated with the spatial pattern of cortical expansion from macaque to human from [128] (each data-point represents one brain region); *Spearman*^*′*^*sr* = *−*0.35, *p* = 0.002 after accounting for spatial autocorrelation.

### Spatial correspondences between macroscopic gradients

So far, we considered all correspondences for an individual gene and protein basis. However, a rich literature posits that much of the spatial variation in gene expression follows a small set of dominant gradients, often identified by applying low-dimensional projections such as principal component analysis [16, 28, 41, 53, 54]. We therefore asked whether the results we observed so far (moderate correspondence between gene expression and protein density in the macaque; greater correspondence between gene expression in the macaque and human) can also be observed at the level of multivariate gradients.

Demonstrating the biological validity of our macaque gene PC1, we show that it is significantly spatially correlated with macaque intracortical myelination (quantified from T1w:T2w ratio), as expected from previous results in both humans and mice [16, 41] (*r* = *−*0.58, *p* = 0.02; Fig. 7a). We replicate this result using a different dataset of macaque intracortical myelination for a subset of regions (Fig. S12). However, corroborating the moderate pair-wise correspondence between macaque genes and receptors, the macaque gene PC1 does not exhibit significant within-species spatial correlation with the macaque receptor PC1 from autoradiography, once spatial autocorrelation is taken into account (*r* = 0.44, *p* = 0.14; Fig. 7b). In contrast, reflecting the inter-species correspondence of gene expression, we find that in addition to the significance of correlations between numerous individual genes across the two species, there is also a significant spatial correlation between the principal component of brain-related gene expression (spatial pattern accounting for most of the variability in the data) in humans and in macaques despite being obtained from different techniques (*r* = 0.64, *p <* 0.001; Fig. 7c).

**Figure 7.**
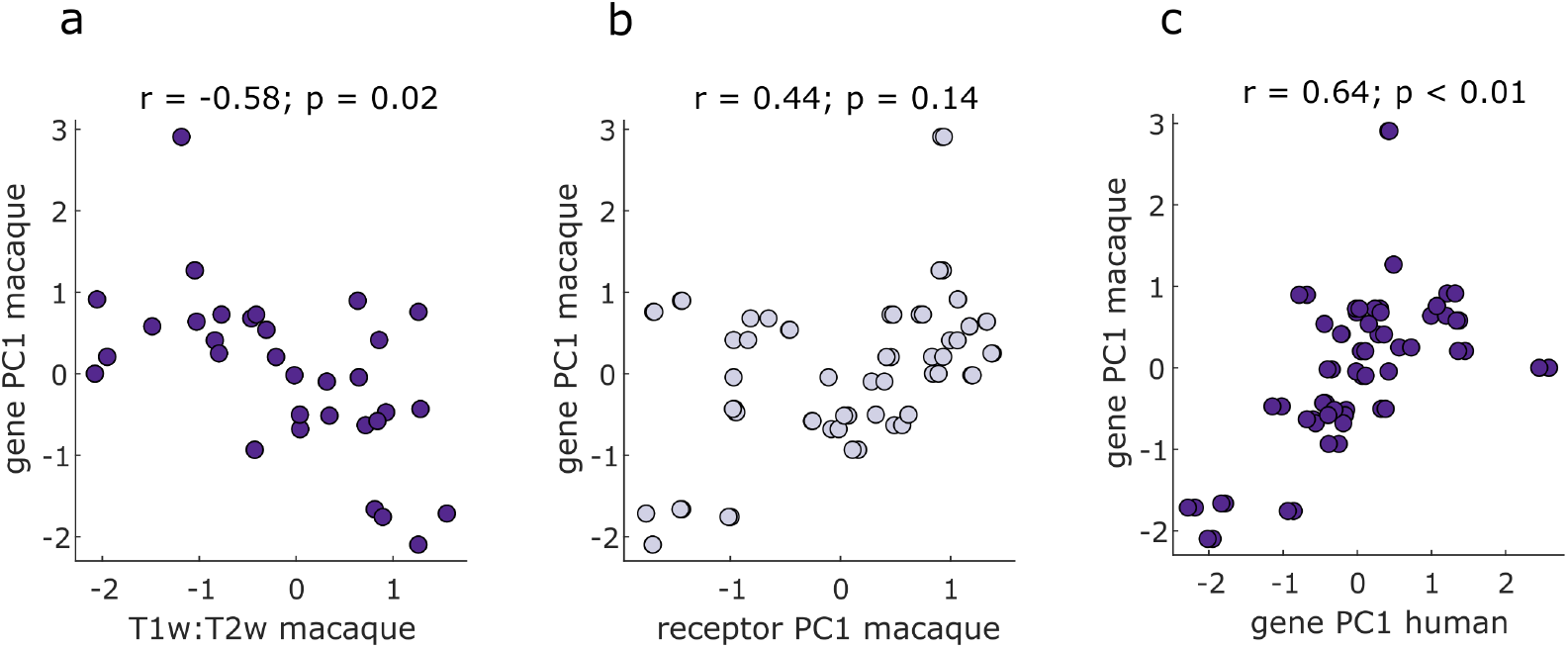
Spatial correspondence across macaque gradients. (**a**) Macaque gene PC1 is significantly spatially correlated with macaque intracortical myelination (quantified from T1w:T2w ratio). (**b**) Macaque gene PC1 does not correlate with macaque receptor PC1. (**c**) Macaque gene PC1 is significantly spatially correlated with human gene PC1. Indigo scatter plots indicate significant spatial correlation (Spearman’s *r, p <* 0.05 against a null distribution with preserved spatial autocorrelation). Values are z-scored.

### Replication with macaque RNA-seq gene expression

We first replicate results using an independent transcriptomics dataset of macaque region-specific bulk RNAsequencing [12]. These data comprise a smaller set of genes coding for neurotransmitter receptors, including some overlapping with the genes used in the present report: *DRD2, GABRQ, GRIA4, GRM4, HTR1B, HTR2C* and *SLC6A2*. We find significant spatial correlation between macaque stereo-seq and RNA-seq for 5*/*7 genes (Fig. S13). Replicating our stereo-seq results for macaque-human correspondence, we find significant spatial correlations between macaque RNA-seq gene expression and human microarray gene expression, for the four genes that were available for both human and macaque (*DRD2, GRIA4, GRM4, HTR2C*) (Fig. S14). Finally, we compare macaque RNA-seq gene expression against macaque receptor density (Fig. S14). We observe moderate correlations between *AMPA* receptor and *GRIA4* gene expression, and between *GABRQ* gene expression and *GABA*_*A*_ receptor density, but neither reached significance in the RNA-seq data (*r* = 0.22, *p* = 0.05 and *r* = 0.42, *p* = 0.06). We do find correspondence between *α*_*2*_ receptor and *ADRA2C* gene expression; although the latter gene was not available in the stereo-seq dataset, we did observe *α*_*2*_ receptor to correlate with *ADRA2A* gene expression (Fig. S15).

### Replication with human RNA-seq gene expression

We also replicate results using an alternative measurement of human gene expression, RNA-seq, available in 2/6 AHBA donors [54]. Patterns of spatial correspondence between human RNA-seq gene expression and macaque stereo-seq gene expression are consistent with microarray results, including significant correlations between human and macaque *PVALB, SST, CALB2, GRIA1, GRIK2* and *HTR1A* (Fig. S16). We also find similar correspondence between human RNA-seq and macaque RNAseq S17. Univariate correspondences are also recapitulated at the multivariate level, where we replicate the significant correlation between macaque gene PC1 and human gene PC1 obtained from RNA-seq (Fig. S18). Lastly, the pattern of inter-species correlation of each region’s gene expression, which we obtained by comparing macaque stereo-seq against human microarray data, is also recapitulated when human RNA-seq data are used instead (Fig. S19). Collectively, these checks demonstrate that our results are robust across different datasets and measurements (human microarray and RNA-seq; macaque stereo-seq and RNA-seq).

## DISCUSSION

Comprehensive gene expression in the macaque holds promise as a means of learning about correspondence with the human brain, as a proxy for other neuroimaging modalities in the macaque, and also to expand the value of the macaque as a model organism, including for gene therapy [37, 60]. Here we sought to comprehensively benchmark the potential for vertical (within species) and horizontal (between species) translation [78] using recently released spatial transcriptomics dataset from *Macaca fascicularis* [12, 22]. We systematically compared the spatial distribution of gene expression in the macaque cortex with (a) macaque cortical receptor density and intracortical myelination and (b) cortical gene expression in the human. As we outline below, we find moderate correspondence between gene expression and receptor density in the macaque, but greater correspondence with other micro-architectural features (calciumbinding proteins, T1w:T2w ratio) and between gene expression in the macaque and human.

Within-species correspondence between macaque gene expression and receptor density is generally moderate, and variable across receptor-gene combinations. This result is in line with the modest correspondence between mRNA and protein abundance previously reported in the literature [6, 14, 27, 49, 99, 118, 134], including in human brain tissue [47, 123]. More importantly, our results validate recent results pertaining specifically to gene-receptor correspondence in the human brain, which showed that relatively few receptors (whether measured from *in vivo* PET or post-mortem autoradiography) exhibit significant spatial covariation with expression patterns of the corresponding genes from human microarray data [50, 63, 64, 81, 136]. Nonetheless, we do find that some specific gene-protein pairs exhibit robust spatial correlations in the macaque brain. These include parvalbumin-*PVALB* and calretinin-*CALB2* correspondence, consistent with the literature in both human and mouse [16, 41], as well as significant spatial correlation between *HTR1A* gene expression and *5HT*_*1A*_ receptor density in the macaque. In the human, [50] showed that the *HTR1A* gene was the only one to show significant correlation with expression of the corresponding receptor (*5HT*_*1A*_) across both PET and autoradiography. Correspondence between *HTR1A* mRNA and (*5HT*_*1A*_) receptor density has been reported in humans by multiple studies [8, 63, 81, 94, 112]. The present results not only corroborate the results obtained by [50, 81, 136] in a different species and with different modalities, but they also validate the recent result of [39], who reported a significant correlation between human *HTR1A* gene expression and macaque *5HT*_*1A*_ receptor density. Likewise, [50] and [81] reported significant correlation between *ADRA2A* and *α*_*2*_ in humans, and [39] reported significant correlation between macaque *α*_*2*_ density and human *ADRA2A* gene expression. Taken together, these previous results suggest that *ADRA2A* expression and *α*_*2*_ density might also be correlated in the macaque—which is what we found. Importantly, although we observed limited gene-receptor correspondence in the macaque when considering gene expression aggregated across cortical layers, 60% of gene-receptor pairs (i.e., twice as many) exhibited a significant correlation for at least one cortical layer. This finding suggests that the limited gene-receptor correspondence reported in the human literature [50, 81] may partly be due to the aggregation of gene expression across cortical layers.

More broadly, there are numerous biological reasons why expression of a gene may diverge from density of the corresponding protein [73, 86]. First, total gene expression level does not take into account the mRNA stability resulting from post-transcriptional regulation, which may influence protein abundance, since RNA abundance of each gene is determined by its rates of transcription and RNA decay [1]. Second, variation in protein stability and thus half-life of proteins as well as the modulated activity of protein transport machinery which determines the final subcellular localization of the protein may affect protein abundance [6, 69, 102, 130]. Rates of degradation in postmortem tissue can vary widely for mRNA and proteins, spanning several orders of magnitude: minutes for mRNA, hours to years for proteins [14, 73, 86, 99, 136]. Third, the proportion of different cell types in a tissue sample may distort the gene expression-protein density correspondence due to differences in the proteome and transcriptome [107, 135]. Fourth, studies in yeast [57] have demonstrated that translation rate also alters protein levels and correlation between mRNA expression and protein abundance. Fifth, the protein itself may be transported far from the coding mRNA, resulting in a spatial discrepancy between mRNA and protein levels—for example in the case of presynaptic receptors located at the end of long axonal projections (note however, that we largely ruled out this possibility by showing that the gene expression does not correlate with the weighted average of a region’s synaptically connected neighbours’ receptor density).

Altogether, the variation in the activity of any of these biological processes may contribute to the observed difference in levels of mRNA expression and protein density in the cortex, including both calcium-binding proteins and receptors. However, since neurotransmitter receptors are protein complexes bound in the cell membrane, there are additional receptor-specific biological processes that may explain why gene-receptor correspondence was generally not as good as the correspondence between gene expression and parvalbumin and calretinin. This is because there are multiple steps between synthesis of a single protein and it being found forming part of a multimeric structure embedded in the cellular membrane. Any formed receptors that do not bind to the membrane will fail to be detected from autoradiography. Additionally, post-transcriptional and post-translational modifications such as receptor assembly and trafficking as well as variations in subunit composition, which play an integral role in synaptic function and plasticity [2, 18, 25, 48, 68, 74, 100, 126], can further contribute to decoupling mRNA expression from the final receptor prevalence.

We find that more than half (51%) of genes pertaining to receptor and neural cell type markers exhibit significant spatial correlation between human and macaque. How the spatial transcriptomes of the macaque and human align across the cortical sheet is an important benchmark for the translational potential of this model organism. Interestingly, we find that inter-species correspondence is regionally heterogeneous but highly systematic. Namely, correspondence is greatest in unimodal cortex and lowest in transmodal cortex – consistent with [7], who reported that similarity of gene expression patterns between human and mouse is higher in sensorimotor than in association cortices. This pattern of genetic divergence may then drive divergence in cell types, circuits, and macroscopic features of cortical organization; for example, we find that regions in which human gene expression diverges most from macaque also show the greatest expansion in grey matter, as reported by Xu and colleagues [128]. This regional heterogeneity bears an important implication for future spatial transcriptomics studies; namely, confidence in translational inferences depends on where in the brain the transcriptome is sampled. Therefore, for horizontal translation from macaque to human it is desirable that transcriptomic sampling should span a variety of brain regions (ideally, whole brain) and be accompanied by precise 3-D spatial locations.

Correspondence between human and macaque is also heterogeneous in terms of specific genes. Once again, however, this heterogeneity is not random. On the contrary, several genes displaying high inter-species correspondence between human and macaque are also known to be conserved between humans and mice. Using each species’ T1w:T2w ratio map to define a common spatial reference for inter-species comparison, previous work identified a number of brain-related genes that exhibit strong inter-species consistency between human and mouse, including the interneuron markers *Pvalb* and *Calb2*, the oxytocin receptor gene *Oxtr*; the glutamate receptor genes *Grin3a, Grik1, Grik2*, and *Grik4*; the myelin marker genes *Mobp* and *Mbp*; and the serotonin *htr1a* gene [41]. Here we find that, except for *Grik4* and *Mbp*, all these genes exhibit significant correlations between human microarray and macaque stereo-seq data, with most (including *GRIK4*) also correlating with human RNA-seq data (note that we used direct spatial correlation between human and macaque cortical expression patterns, rather than the indirect approach of [41], thanks to the availability of the same parcellation in both species). Together with the results of [41], our results further establish that many genes pertaining to microstructure and receptors exhibit robust spatial similarities across mammalian species, even when measured through diverse techniques (in situ hybridization in the mouse; microarray and RNA-seq in humans; stereo-seq in macaques).

This comparative work integrates multiple unique datasets, entailing inherent methodological limitations for each dataset and for comparisons between datasets, including: (1) macaque gene and receptor expression come from different animals; (2) data are post-mortem rather than *in vivo*, and the consequent small number of animals and humans that provided data; (3) gene expression was estimated from different transcriptomics techniques in different species (microarray and RNA-seq for humans; stereo-seq and RNA-seq for macaques); (4) limited cortical coverage for the macaque gene expression data (left hemisphere only); (5) limited cortical coverage for the autoradiography data (no temporal lobe); (6) restriction to cortical sampling.

In light of these limitations, we instituted several checks. We showed that the principal component of macaque gene expression correlates with macaque T1w:T2w ratio map (Fig. 7a), as expected based on analogous correspondences in both human and mouse [16, 41]. Macaque gene PC1 also correlated with human gene PC1, but not with macaque receptor PC1— recapitulating in a multivariate fashion the results observed for individual genes and receptors. We also showed that macaque *PVALB* gene expression correlates with human *PVALB* expression, as expected based on the strong inter-species correspondence between human *PVALB* and mouse *Pvalb* [41]). Using an independent dataset, we further showed that macaque *PVALB* also correlates with parvalbumin protein density in the macaque. Together with the correspondence between *HTR1A* and *5HT*_*1A*_, this evidence indicates that when there is reason from the literature to expect a correspondence between protein density and gene expression, it is also observed in our data. In particular, the correspondence between *HTR1A* and *5HT*_*1A*_ was observed in both supragranular (L2, L3) and infragranular (L5, L6) layers of macaque cortex, consistent with human findings [50], even though our own layer-wise data were for the gene expression, whereas [50] used layer-wise information about receptor density. Notably, *5HT*_*1A*_ receptors are expressed more prominently near the soma (including the axon initial segment), than in the apical dendrite [91], which may provide a neuroanatomical explanation for their higher correlation in layers 2/3/5. In contrast, the high glutamatergic gene-receptor correlations for layer 4 may occur because L4 neurons do not have big dendritic trees (in particular, they do not have apical dendrites). L2/3 and L5 pyramidal cells have broad dendritic trees with apical dendrites reaching up to L1, and therefore can be expected to have many receptors far from the soma. Lastly, we replicated all results using different macaque gene expression data, different protein measurements; and different human gene expression data.

As a final limitation, the datasets used here were each originally provided according to different parcellation schemes and different granularity of sampling. To make all data comparable, we opted to use the canonical Regional Mapping parcellation of the macaque cortex devised by Kötter and Wanke [9, 11, 66], for which tract-tracing and diffusion tractography have already been integrated into a high-quality structural connectome [108]. A human version of this parcellation has been made available, based on functional/cytoarchitectonic landmark-based registration [10, 83]. Crucially, the Regional Mapping atlas was devised with the explicit goal of resolving conflicts between different parcellations in humans and other primate species, by capturing the most consistent cytoarchitectonic, topographic, or functionally defined regions that appear across primate specie s[10, 66] – making it especially well-suited for the needs of the present study to translate between datasets provided in different parcellations. However, alternative macaque atlases exist [113], and cross-species brain mapping is a rapidly-evolving field [32, 35, 55, 71, 124, 128]; different parcellation approaches (e.g., based on different architectonic features, or different imaging modalities) may not always coincide about the exact correspondence between specific cortical locations, especially for densely-sampled data such as functional and structural MRI [55]. Indeed, whereas here we have used a common parcellation to compare different data modalities within macaque and between macaque and human while keeping regional identity fixed, future efforts may take the opposite approach. On one hand, combining some of the recently released datasets about different macaque architectural features is bound to yield richer parcellations of the macaque brain [44, 55]. On the other hand, the availability of gene expression datasets in both humans and macaques may provide the means to enhance existing characterisations of cortical alignment between the two species based on similarity of gene expression, as recently done between human and mouse [7].

Nonetheless, despite the limitations that come with needing to map each original dataset onto a common parcellation, we are reassured that this step has not unduly influenced our results. Our findings are highly consistent with the literature in other species (e.g., correspondence of T1w:T2w ratio with myelin-related genes and *PVALB, GRIN3A*, and gene PC1; *PVALB*-parvalbumin and *CALB2*-calretining correspondence; *HTR1A* and *5HT*_*1A*_), suggesting that the quality of our mapping of the different datasets into a common atlas is sufficient to identify a significant correspondence, where one is expected. Note also that we assessed the statistical significance of each correlation against null distributions of maps with preserved spatial autocorrelation, ensuring that our results are not driven by the role of spatial symmetry or granularity of the data [77]. Lastly, the Regional Mapping atlas is the parcellation scheme used by the popular TheVirtualBrain (TVB) platform for multimodal data sharing and computational modelling [93, 97, 108], enabling smooth integration of macaque gene and receptor expression data with other available data modalities and brain modelling workflows.

In summary, we find moderate within-species gene-receptor correspondence in the macaque cortex, with notable layer-specificity. In contrast, there is better inter-species correspondence for many genes underlying fundamental aspects of brain organisation, such as cell types, receptor subunits, and myelination. This interspecies correspondence of gene expression exhibits systematic regional heterogeneity, giving rise to a cortical pattern of inter-species divergence that recapitulates the inter-species divergence in cortical morphometry. Throughout, our results demonstrate excellent concordance with findings from the translational neuroscience literature in other organisms (similarity of gene expression between human and mouse; limited gene-receptor correspondence in humans) and also using alternative techniques such as immunohistochemistry and T1w:T2w MRI contrast. Taken together, the present results showcase both the potential and limitations of macaque spatial transcriptomics as an engine of translational discovery within and across species.

## METHODS

### Macaque cortical gene expression from stereo-seq

We used cortex-wide macaque gene expression data recently made available by [22], who combined single-nucleus RNA sequencing (“snRNA-seq”) with high-resolution, large-field-of view spatial transcriptomics from spatiotemporal enhanced resolution omics-sequencing (“stereo-seq”) [21]. Specifically, the authors made available (https://macaque.digital-brain.cn/spatial-omics) post-mortem gene expression data covering 143 regions of the left cortical hemisphere of one 6yo male cynomolgus macaque (*Macaca fascicularis*). We refer the reader to [22] for details. The animal protocol was approved by the Biomedical Research Ethics Committee of CAS Center for Excellence in Brain Science and Intelligence Technology, Chinese Academy of Sciences (ION-2019011). Animal care complied with the guideline of this committee [22].

Briefly, Chen and colleagues obtained 119 coronal sections at 500-*µ*m spacing, covering the entire cortex of the left hemisphere, which were used for stereo-seq transcriptomics [22]. Adjacent 50-*µ*m thick sections were also acquired for regional microdissection and snRNAseq analysis, as well as 10-*µ*m sections adjacent to each stereo-seq section, which were used for the anatomical parcellation of brain regions via immunostaining [22]. Stereo-seq is a DNA nanoball (DNB) barcoded solid-phase RNA capture method [21]. It involves reverse transcription of RNAs released from frozen tissue sections fixated onto the stereo-seq chip, and subsequent PCR amplification. The resulting “amplified-barcoded complementary DNA (cDNA) is used as template for library preparation, and sequenced” to obtain high-resolution spatially resolved transcriptomics [21].

Gene expression data were made available for 143 cortical regions of the left hemisphere, including prefrontal, frontal, cingulate, somatosensory, insular, auditory, temporal, parietal, occipital and piriform areas. As reported in [22], for each coronal section, the cortical region and layer parcellation were manually delineated on Stereoseq data background, based on cytoarchitectual pattern (e.g. cell density, cell size) revealed by total mRNA expression, nucleic acid staining, and NeuN staing of adjacent sections. To make the gene expression data comparable across our datasets, the combined gene expression across layers was manually mapped onto the cortical regions of the “regional mapping” macaque atlas of Kötter and Wanke [66], mirroring data between hemispheres.

### Macaque receptor density from *in vitro* receptor autoradiography

In vitro autoradiography data for 14 neurotransmitter receptors were obtained from [39]: *AMPA, kainate, NMDA, GABA*_*A*_, *GABA*_*B*_, *GABA*_*A/BZ*_, *M*_*1*_, *M*_*2*_, *M*_*3*_, *α*_*1*_, *α*_*2*_, *5HT*_*1A*_, *5HT*_*2A*_, *D*_*1*_. The authors applied quantitative in vitro receptor autoradiography to label 14 neurotransmitter receptors in three male *Macaca fascicularis* brains (7.3*±*0.6 years old; body weight 6*±*0.8 kg) obtained from Covance Preclinical Services, where they were housed and used as control animals for pharmaceutical studies performed in compliance with legal requirements. Animal experimental procedures and husbandry had the approval of the respective Institutional Animal Care and Use Committee and were carried out in accordance with the European Council Directive of 2010 [39]. We refer the reader to [39] for details.

Briefly, brain tissue was serially sectioned in the coronal plane (section thickness 20 *µ*m) using a cryostat microtome (Leica, CM3050S). Sections were thaw mounted on gelatine-coated slides, sorted into 22 parallel series and freeze dried overnight. Receptor binding protocols encompass a pre-incubation to rehydrate sections, a main incubation with a tritiated ligand in the presence of or without a non-labeled displacer and a final rinsing step to terminate binding. Incubation with the tritiated ligand alone demonstrates total binding; incubation in combination with the displacer reveals the proportion of non-specific binding sites. Specific binding is the difference between total and non-specific binding and was less than 5% of the total binding. Thus, total binding is considered to be equivalent of specific binding. Sections were exposed together with standards of known radioactivity against tritium-sensitive films (Hyperfilm, Amersham) for 4–18 weeks depending on the receptor type. Ensuing autoradiographs were processed by densitometry with a video-based image analyzing technique. In short, autoradiographs were digitized as 8-bit images. Gray values in the images of the standards were used to compute a calibration curve indicating the relationship between gray values in an autoradiograph and binding site concentrations in femtomole per milligram (fmol *·* mg^*−*1^) of protein. Concentrations of radioactivity (R, counts per minute) in each standard, which had been calibrated against brain tissue homogenate, were converted to binding site concentrations (Cb, fmol *·* mg^*−*1^ of protein). The ensuing calibration curve was used to linearize the autoradiographs—that is, to convert the gray value of each pixel into a binding site concentration in fmol *·* mg^*−*1^ of protein [39].

The data of density of receptors per neuron were made available for 109 cortical areas of the macaque brain, which were identified based on their cytoarchitecture and receptor-architecture characteristics [39]. To enable comparison across datasets, we re-sampled these data to the Regional Mapping macaque cortical parcellation of Kötter and Wanke [66] (see Fig. S20a,b).

### Human gene expression

Regional human gene expression profiles were obtained using microarray data from the Allen Human Brain Atlas (AHBA) [54], with preprocessing as recently described [132]. The Allen Human Brain Atlas (AHBA) is a publicly available transcriptional atlas containing gene expression data measured with DNA microarrays and sampled from hundreds of histologically validated neuroanatomical structures across normal postmortem human brains from six donors (five male and one female; age 24–55 years). We extracted and mapped gene expression data to the 82 cortical ROIs of our parcellation using the *abagen* toolbox https://abagen.readthedocs.io/ [75]. Data were pooled between homologous cortical regions to ensure adequate coverage of both left (data from six donors) and right hemisphere (data from two donors). Distances between samples were evaluated on the cortical surface with a 2mm distance threshold. Only probes where expression measures were above a background threshold in more than 50% of samples were selected. A representative probe for a gene was selected based on highest intensity. Gene expression data were normalised across the cortex using scaled, outlier-robust sigmoid normalisation. 15, 633 genes survived these preprocessing and quality assurance steps. For comparison with the macaque data, the human gene expression data were parcellated into a human-adapted version of the cortical parcellation of Kötter and Wanke [66], as adapted to the human brain by [10] (see Fig. S20a,b). We only included in our analyses genes with a one-toone ortholog between *Homo sapiens* and *Macaca fascicularis*. We also replicate our main results using human gene expression data from an alternative modality, RNAseq, which was available from 2/6 AHBA donors [54].

### Gene-receptor pairs

Neurotransmitter receptors are protein complexes bound in the cell membrane. They can be classified as ionotropic (if signal transduction is mediated by ion channels) or metabotropic (G-protein coupled). Ionotropic receptors are multimeric complexes consisting of multiple subunits, each encoded by a distinct gene. Metabotropic receptors are monomeric complexes: although there are several associated second messenger components, there is only a single protein embedded in the membrane, which is therefore encoded by a single gene. Therefore, for monomeric receptors, we correlated protein density with expression of the corresponding gene. For multimeric receptors, which involve distinct subunits, in our main analysis we correlated protein density with expression of genes coding for the same subunits as used in [50].

- *GABA*_*A*_: this pentamer can be coded by up to 19 subunits. In our main analysis we show correlations with genes coding for the three primary subunits: *α*_1_, *β*_2_ and *γ*_2_.
- *GABA*_*B*_: our main analysis reports correspondence with genes coding for both subunits.
- *AMPA*: in our main analysis we show correlations with the *GRIA1* gene. Correlations with additional genes as used in [81] are reported in the Supplementary.
- *NMDA*: in our main analysis we show correlations with the *GRIN1* gene, which encodes the N1 subunit. Correlations with additional genes as used in [81] are reported in the Supplementary.
- *Kainate*: we show results for the *GRIK2* gene in our main analysis. Correlations with additional genes as used in [81] are reported in the Supplementary.

Since macaque gene expression data from [22] do not include the *CHRM3* gene that codes for the *M*_*3*_ receptor, this receptor could not be included in our analyses. Macaque gene expression was also not available for *ADRA1C* (pertaining to the *α*_*1*_ receptor), so we only included genes *ADRA1A* and *ADRA1B*. Likewise, *ADRA2B* and *ADRA2C* genes (pertaining to the *α*_2_ receptor) were not available, so we only used *ADRA2A*.

### Macaque T1w:T2w maps and parvalbumin and calretinin immunohistochemistry

For our main analysis, we used the map of macaque intracortical myelination from T1w:T2w ratio made available by [39]. These data were available in the same 109area parcellation of the macaque cortex as the receptor data. We therefore followed the same procedure and resampled these data to the Regional Mapping macaque cortical parcellation of Kötter and Wanke [66].

Burt and colleagues [16] assembled data on the mmunohistochemically measured densities of calretinin(also known as calbinding-2) and parvalbumin-expressing inhibitory interneurons for several macaque brain areas, from multiple immunohistochemistry studies [26, 31, 42, 65]. The same authors also provide T1w:T2w intracortical myelination data for the same regions [16], which we used for our replication analysis. To compare data across modalities, we manually mapped their data onto 38 bilateral regions of the Regional Mapping parcellation [66].

In addition to the aggregated data of Burt and colleagues [16], we also used immunohistochemistry data about the prevalence of parvalbumin-immunoreactive and calretinin-immunoreactive neurons for a subset of macaque visual, auditory, and somatosensory cortical regions from [65], which was one of the original studies combined by [16]. Data were obtained from three normal adult macaque monkeys (*Macaca fuscata*), with approval from the animal research committee of RIKEN (Japan) and in accordance with the Guiding Principles for the Care and Use of Animals in the Field of Physiological Sciences of the Japanese Physiological Society [65]. Briefly, a monoclonal antiparvalbumin antibody (Sigma) and a polyclonal anticalretinin antiserum (SWant) were used on post-mortem brain sections (30 microns of thickness). The number of parvalbumin-immunoreactive and calretinin-immunoreactive neurons (all non-pyramidal except for a few calretinin-immunoreactive pyramidal neurons in area 28) was counted in 200-micron-wide strips extending vertically to the cortical surface through all layers, and for each cortical area this process was repeated 16 times at different locations in the three monkeys [65]. Here, we ranked areas based on their mean count of immunoreactive neurons (separately for calretinin and parvalbumin) as displayed in Figure 4 of [65], and subsequently manually mapped these ranks onto bilateral regions of the Regional Mapping parcellation [66].

### Brain-related genes

In addition to the lists of receptor-related genes from [50, 81], we also obtained from [41] a list of 124 brain-related genes coding for receptor subunits, as well as cell-type markers for parvalbumin, somatostatin, vasoactive intestinal peptide, calbindin, and myelin. We further added genes coding for SLC6 transporters, histidine decarboxylase (*HDC*), and syntaxin (*STX1A*), hyperpolarization-activated cyclic nucleotide-gated channels (HCN), potassium channels (KCN), and sodium voltage-gated channels (SCN), which are important for brain function. Of these, 106 genes were available and passed our quality control in both macaque and human, and were included in our analysis.

### Alternative macaque gene expression data from RNA-seq

Bo et al. [12] made available bulk RNA-sequencing regional expression patterns of several neurotransmitter-related genes from the brain of *Macaca fascicularis*, including some overlapping with the list of brain-related genes used in the present study: *DRD2, GABRQ, GRIA4, GRM4, HTR1B, HTR2C* and *SLC6A2*. Expression patterns are provided for 97 regions of the D99 macaque atlas, and we manually mapped them onto the Regional Mapping parcellation [66]. As reported in the original study [12], 9 adult cynomolgus monkeys (*Macaca fascicularis*; mean *±* s.d., 13.6 *±* 7.8 years, 8 males and 1 female) weighing 4.2–12.0 kg (8.6 *±* 2.6 kg) were used for the study. All animal experimental procedures were approved by the Animal Care and Use Committee of CAS Center for Excellence in Brain Science and Intelligence Technology, Chinese Academy of Sciences, and conformed to National Institutes of Health guidelines for the humane care and use of laboratory animals. We refer the reader to the original study [12] for details of regional bulk RNA-sequencing procedures.

### Macaque anatomical connectivity

Anatomical connectivity for the macaque brain was obtained from the fully weighted, whole-cortex macaque connectome recently developed by Shen and colleagues [108]. This connectome was generated by combining information from two different axonal tract-tracing studies from the CoCoMac database (http://cocomac.g-node.org/main/index.php) [3, 111] with diffusion-based tractography obtained from nine adult macaques (*Macaca mulatta* and *Macaca fascicularis*) [108]. The resulting connectome provides a matrix of weighted, directed anatomical connectivity between each of the cortical ROIs of the Regional Mapping atlas of Kötter and Wanke [108].

### Macaque-human cortical expansion

To contextualise our regional pattern of inter-species correlation of gene expression, we used *neuromaps* (https://netneurolab.github.io/neuromaps/) [76] to obtain the map of cortical expansion between macaque and human from [128], parcellated into Schaefer-400 cortical atlas [98].

### Statistical analyses

Spatial correspondence across cortical regions was quantified using Spearman’s rank-based correlation coefficient, which is more robust to outliers and nonnormality than Pearson correlation, and is recommended when studying the correlation between mRNA and protein levels [73]. To control for the spatial autocorrelation inherent in neuroimaging data, which can induce an inflated rate of false positives [77, 115], we assessed the statistical significance of correlations non-parametrically, by comparing each empirical correlation against a distribution of 10, 000 correlations with null maps having the same spatial autocorrelation. Null maps were generated using Moran spectral randomisation on the inverse Euclidean distances between parcel centroids, as implemented in the *BrainSpace* toolbox (https://brainspace.readthedocs.io/en/latest/) [119]. Moran spectral randomisation quantifies the spatial autocorrelation in the data in terms of Moran’s *I* coefficient [24, 33, 120], by computing spatial eigenvectors known as Moran eigenvector maps. The Moran eigenvectors are then used to generate null maps data by imposing the spatial structure of the empirical data on randomised surrogate data [77, 119]. Comparisons between proportions of significantly correlated patterns (between human and macaque genes, or macaque genes and receptors) were performed using the *χ*^2^ test.

## Data and code availability

The *abagen* toolbox for processing of the AHBA human transcriptomic dataset is available at https://abagen.readthedocs.io/. The *neuromaps* toolbox is available at https://netneurolab.github.io/neuromaps/. The *BrainSpace* toolbox is available at https://brainspace.readthedocs.io/en/latest/. Macaque cortical gene expression data from [22] are available at https://macaque.digital-brain.cn/spatial-omics. The dataset is provided by Brain Science Data Center, Chinese Academy of Sciences (https://braindatacenter.cn/). Macaque receptor density data from autoradiography are available from https://balsa.wustl.edu/study/P2Nql and https://search.kg.ebrains.eu/instances/de62abc1-7252-4774-9965-5040f5e8fb6b [38, 39]. The map of macaque intracortical myelination from T1w:T2w ratio from [39] is available at https://balsa.wustl.edu/study/P2Nql. Macaque parvalbumin and calretinin density from immunohistochemistry are available from the Supplementary Materials of [16].

## Acknowledgments

We thank Gleb Bezgin for developing and sharing the human version of the Regional Mapping atlas. We also thank members of the Network Neuroscience Lab at McGill University for valuable comments. We acknowledge the support of the Helmholtz Association’s Initiative and Networking Fund through the Helmholtz International BigBrain Analytics and Learning Laboratory (HIBALL) under the Helmholtz International Lab (grant agreement InterLabs-0015); AIL acknowledges support from the Natural Sciences and Engineering Research Council of Canada (NSERC), [funding reference number 202209BPF-489453-401636, Banting Postdoctoral Fellowship] and FRQNT Strategic Clusters Program (2020RS4-265502 Centre UNIQUE Union Neuroscience & Artificial Intelligence Quebec) via the UNIQUE NeuroAI Excellence Award. BM acknowledges support from the Natural Sciences and Engineering Research Council of Canada (NSERC), Canadian Institutes of Health Research (CIHR), Brain Canada Foundation Future Leaders Fund, the Canada Research Chairs Program, the Michael J. Fox Foundation, and the Healthy Brains for Healthy Lives initiative. ZQL acknowledges support from the Fonds de Recherche du Québec – Nature et Technologies (FRQNT). Any opinions, findings, and conclusions or recommendations expressed in this material are those of the authors and do not reflect the views of the funders.

## References

[1] Alkallas, R., Fish, L., Goodarzi, H., and Najafabadi, H. S. (2017). Inference of rna decay rate from transcriptional profiling highlights the regulatory programs of alzheimer’s disease. Nature communications, 8(1):909.

[2] Anne Stephenson, F., Cousins, S. L., and Kenny, A. V. (2008). Assembly and forward trafficking of nmda receptors. Molecular membrane biology, 25(4):311–320.

[3] Bakker, R., Wachtler, T., and Diesmann, M. (2012). Cocomac 2.0 and the future of tract-tracing databases. Frontiers in neuroinformatics, 6:30.

[4] Ballentine, G., Friedman, S. F., and Bzdok, D. (2022). Trips and neurotransmitters: Discovering principled patterns across 6850 hallucinogenic experiences. Science advances, 8(11):eabl6989.

[5] Barron, H. C., Mars, R. B., Dupret, D., Lerch, J. P., and Sampaio-Baptista, C. (2021). Crossspecies neuroscience: closing the explanatory gap. Philosophical Transactions of the Royal Society B, 376(1815):20190633.

[6] Battle, A., Khan, Z., Wang, S. H., Mitrano, A., Ford, M. J., Pritchard, J. K., and Gilad, Y. (2015). Impact of regulatory variation from rna to protein. Science, 347(6222):664–667.

[7] Beauchamp, A., Yee, Y., Darwin, B. C., Raznahan, A., Mars, R. B., and Lerch, J. P. (2022). Whole-brain comparison of rodent and human brains using spatial transcriptomics. Elife, 11:e79418.

[8] Beliveau, V., Ganz, M., Feng, L., Ozenne, B., Højgaard, L., Fisher, P. M., Svarer, C., Greve, D. N., and Knudsen, G. M. (2017). A high-resolution in vivo atlas of the human brain’s serotonin system. Journal of Neuroscience, 37(1):120–128.

[9] Bezgin, G., Reid, A. T., Schubert, D., and Kötter, R. (2009). Matching spatial with ontological brain regions using java tools for visualization, database access, and integrated data analysis. Neuroinformatics, 7:7–22.

[10] Bezgin, G., Solodkin, A., Bakker, R., Ritter, P., and McIntosh, A. R. (2017). Mapping complementary features of cross-species structural connectivity to construct realistic “virtual brains”. Human Brain Mapping, 38(4):2080– 2093.

[11] Bezgin, G., Vakorin, V. A., van Opstal, A. J., McIntosh, A. R., and Bakker, R. (2012). Hundreds of brain maps in one atlas: registering coordinate-independent primate neuro-anatomical data to a standard brain. Neuroimage, 62(1):67–76.

[12] Bo, T., Li, J., Hu, G., Zhang, G., Wang, W., Lv, Q., Zhao, S., Ma, J., Qin, M., Yao, X., et al. (2023). Brain-wide and cell-specific transcriptomic insights into mri-derived cortical morphology in macaque monkeys. Nature Communications, 14(1):1499.

[13] Braun, U., Harneit, A., Pergola, G., Menara, T., Schäfer, A., Betzel, R. F., Zang, Z., Schweiger, J. I., Zhang, X., Schwarz, K., et al. (2021). Brain network dynamics during working memory are modulated by dopamine and diminished in schizophrenia. Nature Communications, 12(1):1–11.

[14] Buccitelli, C. and Selbach, M. (2020). mrnas, proteins and the emerging principles of gene expression control. Nature Reviews Genetics, 21(10):630–644.

[15] Buffalo, E. A., Movshon, J. A., and Wurtz, R. H. (2019). From basic brain research to treating human brain disorders. Proceedings of the National Academy of Sciences, 116(52):26167–26172.

[16] Burt, J. B., Demirtaş, M., Eckner, W. J., Navejar, N. M., Ji, J. L., Martin, W. J., Bernacchia, A., Anticevic, A., and Murray, J. D. (2018). Hierarchy of transcriptomic specialization across human cortex captured by structural neuroimaging topography. Nature neuroscience, 21(9):1251–1259.

[17] Burt, J. B., Preller, K. H., Demirtas, M., Ji, J. L., Krystal, J. H., Vollenweider, F. X., Anticevic, A., and Murray, J. D. (2021). Transcriptomics-informed large-scale cortical model captures topography of pharmacological neuroimaging effects of lsd. Elife, 10:e69320.

[18] Campbell, B. F. and Tyagarajan, S. K. (2019). Cellular mechanisms contributing to the functional heterogeneity of gabaergic synapses. Frontiers in molecular neuroscience, 12:187.

[19] Capitanio, J. P. and Emborg, M. E. (2008). Contributions of non-human primates to neuroscience research. The Lancet, 371(9618):1126–1135.

[20] Chang, T.-H., Gloria, Y. C., Hellmann, M. J., Greve, C. L., Le Roy, D., Roger, T., Kasper, L., Hube, B., Pusch, S., Gow, N., et al. (2022). Transkingdom mechanism of mamp generation by chitotriosidase (chit1) feeds oligomeric chitin from fungal pathogens and allergens into tlr2-mediated innate immune sensing. bioRxiv.

[21] Chen, A., Liao, S., Cheng, M., Ma, K., Wu, L., Lai, Y., Qiu, X., Yang, J., Xu, J., Hao, S., et al. (2022). Spatiotemporal transcriptomic atlas of mouse organogenesis using dna nanoball-patterned arrays. Cell, 185(10):1777–1792.

[22] Chen, A., Sun, Y., Lei, Y., Li, C., Liao, S., Meng, J., Bai, Y., Liu, Z., Liang, Z., Zhu, Z., et al. (2023). Single-cell spatial transcriptome reveals cell-type organization in the macaque cortex. Cell, 186(17):3726–3743.

[23] Chiou, K. L., Huang, X., Bohlen, M. O., Tremblay, S., DeCasien, A. R., O’Day, D. R., Spurrell, C. H., Gogate, A. A., Zintel, T. M., Unit, C. B. R., et al. (2023). A single-cell multi-omic atlas spanning the adult rhesus macaque brain. Science Advances, 9(41):eadh1914.

[24] Cliff, A. D. and Ord, J. K. (1973). Spatial autocorrelation. (No Title).

[25] Collingridge, G. L., Isaac, J. T., and Wang, Y. T. (2004). Receptor trafficking and synaptic plasticity. Nature reviews neuroscience, 5(12):952–962.

[26] Condé, F., Lund, J. S., Jacobowitz, D. M., Baimbridge, K. G., and Lewis, D. A. (1994). Local circuit neurons immunoreactive for calretinin, calbindin d-28k or parvalbumin in monkey prefronatal cortex: Distribution and morphology. Journal of Comparative Neurology, 341(1):95–116.

[27] de Sousa Abreu, R., Penalva, L. O., Marcotte, E. M., and Vogel, C. (2009). Global signatures of protein and mrna expression levels. Molecular BioSystems, 5(12):1512– 1526.

[28] Dear, R., Wagstyl, K., Seidlitz, J., Markello, R. D., Arnatkevičiūtė, A., Anderson, K. M., Bethlehem, R. A., Consortium, L. B. C., Raznahan, A., Bullmore, E. T., et al. (2024). Cortical gene expression architecture links healthy neurodevelopment to the imaging, transcriptomics and genetics of autism and schizophrenia. Nature Neuroscience, pages 1–12.

[29] Deco, G., Kringelbach, M. L., Arnatkeviciute, A., Oldham, S., Sabaroedin, K., Rogasch, N. C., Aquino, K. M., and Fornito, A. (2021). Dynamical consequences of regional heterogeneity in the brain’s transcriptional landscape. Science Advances, 7(29):eabf4752.

[30] Diez, I. and Sepulcre, J. (2018). Neurogenetic profiles delineate large-scale connectivity dynamics of the human brain. Nature communications, 9(1):3876.

[31] Dombrowski, S., Hilgetag, C., and Barbas, H. (2001). Quantitative architecture distinguishes prefrontal cortical systems in the rhesus monkey. Cerebral cortex, 11(10):975–988.

[32] Donahue, C. J., Glasser, M. F., Preuss, T. M., Rilling, J. K., and Van Essen, D. C. (2018). Quantitative assessment of prefrontal cortex in humans relative to nonhuman primates. Proceedings of the National Academy of Sciences, 115(22):E5183–E5192.

[33] Dray, S. (2011). A new perspective about moran’s coefficient: Spatial autocorrelation as a linear regression problem. Geographical Analysis, 43(2):127–141.

[34] Ebeling, M., Küng, E., See, A., Broger, C., Steiner, G., Berrera, M., Heckel, T., Iniguez, L., Albert, T., Schmucki, R., et al. (2011). Genome-based analysis of the nonhuman primate macaca fascicularis as a model for drug safety assessment. Genome research, 21(10):1746–1756.

[35] Eichert, N., Robinson, E. C., Bryant, K. L., Jbabdi, S., Jenkinson, M., Li, L., Krug, K., Watkins, K. E., and Mars, R. B. (2020). Cross-species cortical alignment identifies different types of anatomical reorganization in the primate temporal lobe. Elife, 9:e53232.

[36] Feng, G., Jensen, F. E., Greely, H. T., Okano, H., Treue S., Roberts, A. C., Fox, J. G., Caddick, S., Poo, M.-m., Newsome, W. T., et al. (2020). Opportunities and limitations of genetically modified nonhuman primate models for neuroscience research. Proceedings of the National Academy of Sciences, 117(39):24022–24031.

[37] Ford, M. M., George, B. E., Van Laar, V. S., Holleran, K. M., Naidoo, J., Hadaczek, P., Vanderhooft, L. E., Peck, E. G., Dawes, M. H., Ohno, K., et al. (2023). Gdnf gene therapy for alcohol use disorder in male non-human primates. Nature medicine, 29(8):2030–2040.

[38] Froudist-Walsh, S., Rapan, L., Palomero-Gallagher, N., and Niu, M. (2023a). Neurotransmitter receptor densities per neuron across macaque cortex (v1. 0). Technical report, Strukturelle und funktionelle Organisation des Gehirns.

[39] Froudist-Walsh, S., Xu, T., Niu, M., Rapan, L., Zhao, L., Margulies, D. S., Zilles, K., Wang, X.-J., and Palomero-Gallagher, N. (2023b). Gradients of neurotransmitter receptor expression in the macaque cortex. Nature neuroscience, 26(7):1281–1294.

[40] Fulcher, B. D. and Fornito, A. (2016). A transcriptional signature of hub connectivity in the mouse connectome. Proc Natl Acad Sci USA, 113(5):1435–1440.

[41] Fulcher, B. D., Murray, J. D., Zerbi, V., and Wang, X.-J. (2019). Multimodal gradients across mouse cortex. Proc Natl Acad Sci USA, 116(10):4689–4695.

[42] Gabbott, P. L. and Bacon, S. J. (1996). Local circuit neurons in the medial prefrontal cortex (areas 24a, b, c, 25 and 32) in the monkey: Ii. quantitative areal and laminar distributions. Journal of Comparative Neurology, 364(4):609–636.

[43] Gao, R., van den Brink, R. L., Pfeffer, T., and Voytek, B. (2020). Neuronal timescales are functionally dynamic and shaped by cortical microarchitecture. Elife, 9:e61277.

[44] Glasser, M. F., Coalson, T. S., Robinson, E. C., Hacker, C. D., Harwell, J., Yacoub, E., Ugurbil, K., Andersson, J., Beckmann, C. F., Jenkinson, M., et al. (2016). A multimodal parcellation of human cerebral cortex. Nature, 536(7615):171–178.

[45] Glasser, M. F., Goyal, M. S., Preuss, T. M., Raichle, M. E., and Van Essen, D. C. (2014). Trends and properties of human cerebral cortex: correlations with cortical myelin content. Neuroimage, 93:165–175.

[46] Glasser, M. F. and Van Essen, D. C. (2011). Mapping human cortical areas in vivo based on myelin content as revealed by t1-and t2-weighted mri. Journal of neuroscience, 31(32):11597–11616.

[47] Godbersen, G., Murgaš, M., Gryglewski, G., Klöbl, M., Unterholzner, J., Rischka, L., Spies, M., Baldinger-Melich, P., Winkler, D., and Lanzenberger, R. (2022). Coexpression of gene transcripts with monoamine oxidase a quantified by human in vivo positron emission tomography. Cerebral Cortex, 32(16):3516–3524.

[48] Groc, L. and Choquet, D. (2020). Linking glutamate receptor movements and synapse function. Science, 368(6496):eaay4631.

[49] Gry, M., Rimini, R., Strömberg, S., Asplund, A., Pontén, F., Uhlén, M., and Nilsson, P. (2009). Correlations between rna and protein expression profiles in 23 human cell lines. BMC genomics, 10:1–14.

[50] Hansen, J. Y., Markello, R. D., Tuominen, L., Nørgaard, M., Kuzmin, E., Palomero-Gallagher, N., Dagher, A., and Misic, B. (2022a). Correspondence between gene expression and neurotransmitter receptor and transporter density in the human brain. NeuroImage, 264:119671.

[51] Hansen, J. Y., Markello, R. D., Vogel, J. W., Seidlitz, J., Bzdok, D., and Misic, B. (2021). Mapping gene transcription and neurocognition across human neocortex. Nature Human Behaviour, 5(9):1240–1250.

[52] Hansen, J. Y., Shafiei, G., Vogel, J. W., Smart, K., Bearden, C. E., Hoogman, M., Franke, B., Van Rooij, D., Buitelaar, J., McDonald, C. R., et al. (2022b). Local molecular and global connectomic contributions to cross-disorder cortical abnormalities. Nature communications, 13(1):1–17.

[53] Hawrylycz, M., Miller, J. A., Menon, V., Feng, D., Dolbeare, T., Guillozet-Bongaarts, A. L., Jegga, A. G., Aronow, B. J., Lee, C.-K., Bernard, A., et al. (2015). Canonical genetic signatures of the adult human brain. Nat Neurosci, 18(12):1832.

[54] Hawrylycz, M. J., Lein, E. S., Guillozet-Bongaarts, A. L., Shen, E. H., Ng, L., Miller, J. A., Van De Lagemaat, L. N., Smith, K. A., Ebbert, A., Riley, Z. L., et al. (2012). An anatomically comprehensive atlas of the adult human brain transcriptome. Nature, 489(7416):391.

[55] Hayashi, T., Hou, Y., Glasser, M. F., Autio, J. A., Knoblauch, K., Inoue-Murayama, M., Coalson, T., Yacoub, E., Smith, S., Kennedy, H., et al. (2021). The nonhuman primate neuroimaging and neuroanatomy project. Neuroimage, 229:117726.

[56] Hill, J., Inder, T., Neil, J., Dierker, D., Harwell, J., and Van Essen, D. (2010). Similar patterns of cortical expansion during human development and evolution. Proceedings of the National Academy of Sciences, 107(29):13135–13140.

[57] Ho, B., Baryshnikova, A., and Brown, G. W. (2018). Unification of protein abundance datasets yields a quantitative saccharomyces cerevisiae proteome. Cell systems, 6(2):192–205.

[58] Hoftman, G. D., Dienel, S. J., Bazmi, H. H., Zhang, Y., Chen, K., and Lewis, D. A. (2018). Altered gradients of glutamate and gamma-aminobutyric acid transcripts in the cortical visuospatial working memory network in schizophrenia. Biological psychiatry, 83(8):670–679.

[59] Huntenburg, J. M., Bazin, P.-L., and Margulies, D. S. (2018). Large-scale gradients in human cortical organization. Trends Cogn Sci, 22(1):21–31.

[60] Jarraya, B., Boulet, S., Scott Ralph, G., Jan, C., Bonvento, G., Azzouz, M., Miskin, J. E., Shin, M., Delzescaux, T., Drouot, X., et al. (2009). Dopamine gene therapy for parkinson’s disease in a nonhuman primate without associated dyskinesia. Science translational medicine, 1(2):2ra4–2ra4.

[61] Jennings, C. G., Landman, R., Zhou, Y., Sharma, J., Hyman, J., Movshon, J. A., Qiu, Z., Roberts, A. C., Roe, A. W., Wang, X., et al. (2016). Opportunities and challenges in modeling human brain disorders in transgenic primates. Nature Neuroscience, 19(9):1123–1130.

[62] Joyce, M. K. P., Ivanov, T. G., Krienen, F., Mitchell, J., Ma, S., Inoue, W., Nandy, A. P., Datta, D., Duque, A., Arellano, J. I., et al. (2024). Dopamine d1 receptor expression in dlpfc inhibitory parvalbumin neurons may contribute to higher visuospatial distractibility in marmosets versus macaques. bioRxiv, pages 2024–06.

[63] Komorowski, A., James, G., Philippe, C., Gryglewski, G., Bauer, A., Hienert, M., Spies, M., Kautzky, A., Vanicek, T., Hahn, A., et al. (2017). Association of protein distribution and gene expression revealed by pet and postmortem quantification in the serotonergic system of the human brain. Cerebral Cortex, 27(1):117–130.

[64] Komorowski, A., Weidenauer, A., Murgaš, M., Sauerzopf, U., Wadsak, W., Mitterhauser, M., Bauer, M., Hacker, M., Praschak-Rieder, N., Kasper, S., et al. (2020). Association of dopamine d2/3 receptor binding potential measured using pet and [11c]-(+)-phno with post-mortem drd2/3 gene expression in the human brain. Neuroimage, 223:117270.

[65] Kondo, H., Tanaka, K., Hashikawa, T., and Jones, E. G. (1999). Neurochemical gradients along monkey sensory cortical pathways: calbindin-immunoreactive pyramidal neurons in layers ii and iii. European Journal of Neuroscience, 11(12):4197–4203.

[66] Kötter, R. and Wanke, E. (2005). Mapping brains without coordinates. Philosophical Transactions of the Royal Society B: Biological Sciences, 360(1456):751–766.

[67] Lein, E. S., Hawrylycz, M. J., Ao, N., Ayres, M., Bensinger, A., Bernard, A., Boe, A. F., Boguski, M. S., Brockway, K. S., Byrnes, E. J., et al. (2007). Genome-wide atlas of gene expression in the adult mouse brain. Nature, 445(7124):168–176.

[68] Levitz, J., Habrian, C., Bharill, S., Fu, Z., Vafabakhsh, R., and Isacoff, E. Y. (2016). Mechanism of assembly and cooperativity of homomeric and heteromeric metabotropic glutamate receptors. Neuron, 92(1):143– 159.

[69] Liu, Y., Beyer, A., and Aebersold, R. (2016a). On the dependency of cellular protein levels on mrna abundance. Cell, 165(3):535–550.

[70] Liu, Z., Li, X., Zhang, J.-T., Cai, Y.-J., Cheng, T.-L., Cheng, C., Wang, Y., Zhang, C.-C., Nie, Y.-H., Chen, Z.-F., et al. (2016b). Autism-like behaviours and germline transmission in transgenic monkeys overexpressing mecp2. Nature, 530(7588):98–102.

[71] Lu, Y., Cui, Y., Cao, L., Dong, Z., Cheng, L., Wu, W., Wang, C., Liu, X., Liu, Y., Zhang, B., et al. (2024). Macaque brainnetome atlas: A multifaceted brain map with parcellation, connection, and histology. Science Bulletin.

[72] Luppi, A. I., Mediano, P. A. M., Rosas, F. E., Holland, N., Fryer, T. D., O’Brien, J. T., Rowe, J. B., Menon, D. K., Bor, D., and Stamatakis, E. A. (2022). A synergistic core for human brain evolution and cognition. Nature Neuroscience, 25(6):771–782. number: 6 publisher: Nature Publishing Group.

[73] Maier, T., Güell, M., and Serrano, L. (2009). Correlation of mrna and protein in complex biological samples. FEBS letters, 583(24):3966–3973.

[74] Malinow, R. and Malenka, R. C. (2002). Ampa receptor trafficking and synaptic plasticity. Annual review of neuroscience, 25(1):103–126.

[75] Markello, R. D., Arnatkeviciute, A., Poline, J.-B., Fulcher, B. D., Fornito, A., and Misic, B. (2021). Standardizing workflows in imaging transcriptomics with the abagen toolbox. Elife, 10:e72129.

[76] Markello, R. D., Hansen, J. Y., Liu, Z.-Q., Bazinet, V., Shafiei, G., Suárez, L. E., Blostein, N., Seidlitz, J., Baillet, S., Satterthwaite, T. D., et al. (2022). Neuromaps: structural and functional interpretation of brain maps. Nature Methods, 19(11):1472–1479.

[77] Markello, R. D. and Misic, B. (2021). Comparing spatial null models for brain maps. NeuroImage, page 118052.

[78] Mars, R. B., Jbabdi, S., and Rushworth, M. F. (2021). A common space approach to comparative neuroscience. Annual Review of Neuroscience, 44:69–86.

[79] Milham, M., Petkov, C. I., Margulies, D. S., Schroeder, C. E., Basso, M. A., Belin, P., Fair, D. A., Fox, A., Kastner, S., Mars, R. B., et al. (2020). Accelerating the evolution of nonhuman primate neuroimaging. Neuron, 105(4):600–603.

[80] Milham, M. P., Ai, L., Koo, B., Xu, T., Amiez, C., Balezeau, F., Baxter, M. G., Blezer, E. L., Brochier, T., Chen, A., et al. (2018). An open resource for non-human primate imaging. Neuron, 100(1):61–74.

[81] Murgaš, M., Michenthaler, P., Reed, M. B., Gryglewski, G., and Lanzenberger, R. (2022). Correlation of receptor density and mrna expression patterns in the human cerebral cortex. NeuroImage, 256:119214.

[82] Nørgaard, M., Beliveau, V., Ganz, M., Svarer, C., Pinborg, L. H., Keller, S. H., Jensen, P. S., Greve, D. N., and Knudsen, G. M. (2021). A high-resolution in vivo atlas of the human brain’s benzodiazepine binding site of gabaa receptors. NeuroImage, 232:117878.

[83] Orban, G. A., Van Essen, D., and Vanduffel, W. (2004). Comparative mapping of higher visual areas in monkeys and humans. Trends in cognitive sciences, 8(7):315–324.

[84] Pagani, M., Gutierrez-Barragan, D., de Guzman, A. E., Xu, T., and Gozzi, A. (2023). Mapping and comparing fmri connectivity networks across species. Communications Biology, 6(1):1238.

[85] Passingham, R. (2009). How good is the macaque monkey model of the human brain? Current opinion in neurobiology, 19(1):6–11.

[86] Payne, S. H. (2015). The utility of protein and mrna correlation. Trends in biochemical sciences, 40(1):1–3.

[87] Pecheva, D., Lee, A., Poh, J. S., Chong, Y.-S., Shek, L. P., Gluckman, P. D., Meaney, M. J., Fortier, M. V., and Qiu, A. (2020). Neural transcription correlates of multimodal cortical phenotypes during development. Cerebral Cortex, 30(5):2740–2754.

[88] Pembroke, W. G., Hartl, C. L., and Geschwind, D. H. (2021). Evolutionary conservation and divergence of the human brain transcriptome. Genome biology, 22(1):1– 33.

[89] Phillips, K. A., Bales, K. L., Capitanio, J. P., Conley, A., Czoty, P. W., ‘t Hart, B. A., Hopkins, W. D., Hu, S.-L., Miller, L. A., Nader, M. A., et al. (2014). Why primate models matter. American journal of primatology, 76(9):801–827.

[90] Preller, K. H., Burt, J. B., Ji, J. L., Schleifer, C. H., Adkinson, B. D., Stämpfli, P., Seifritz, E., Repovs, G., Krystal, J. H., Murray, J. D., et al. (2018). Changes in global and thalamic brain connectivity in lsd-induced altered states of consciousness are attributable to the 5-ht2a receptor. Elife, 7:e35082.

[91] Puig, M. V. and Gulledge, A. T. (2011). Serotonin and prefrontal cortex function: neurons, networks, and circuits. Molecular neurobiology, 44:449–464.

[92] Richiardi, J., Altmann, A., Milazzo, A.-C., Chang, C., Chakravarty, M. M., Banaschewski, T., Barker, G. J., Bokde, A. L., Bromberg, U., Büchel, C., et al. (2015). Correlated gene expression supports synchronous activity in brain networks. Science, 348(6240):1241–1244.

[93] Ritter, P., Schirner, M., McIntosh, A. R., and Jirsa, V. K. (2013). The virtual brain integrates computational modeling and multimodal neuroimaging. Brain connectivity, 3(2):121–145.

[94] Rizzo, G., Veronese, M., Heckemann, R. A., Selvaraj, S., Howes, O. D., Hammers, A., Turkheimer, F. E., and Bertoldo, A. (2014). The predictive power of brain mrna mappings for in vivo protein density: a positron emission tomography correlation study. Journal of Cerebral Blood Flow & Metabolism, 34(5):827–835.

[95] Roelfsema, P. R. and Treue, S. (2014). Basic neuroscience research with nonhuman primates: a small but indispensable component of biomedical research. Neuron, 82(6):1200–1204.

[96] Romero-Garcia, R., Whitaker, K. J., Váša, F., Seidlitz, J., Shinn, M., Fonagy, P., Dolan, R. J., Jones, P. B., Goodyer, I. M., Bullmore, E. T., et al. (2018). Structural covariance networks are coupled to expression of genes enriched in supragranular layers of the human cortex. NeuroImage, 171:256–267.

[97] Sanz Leon, P., Knock, S. A., Woodman, M. M., Domide, L., Mersmann, J., McIntosh, A. R., and Jirsa, V. (2013). The virtual brain: a simulator of primate brain network dynamics. Frontiers in neuroinformatics, 7:10.

[98] Schaefer, A., Kong, R., Gordon, E. M., Laumann, T. O., Zuo, X.-N., Holmes, A. J., Eickhoff, S. B., and Yeo, B. T. (2018). Local-global parcellation of the human cerebral cortex from intrinsic functional connectivity mri. Cerebral cortex, 28(9):3095–3114.

[99] Schwanhäusser, B., Busse, D., Li, N., Dittmar, G., Schuchhardt, J., Wolf, J., Chen, W., and Selbach, M. (2011). Global quantification of mammalian gene expression control. Nature, 473(7347):337–342.

[100] Schwappach, B. (2008). An overview of trafficking and assembly of neurotransmitter receptors and ion channels. Molecular membrane biology, 25(4):270–278.

[101] Seidlitz, J., Nadig, A., Liu, S., Bethlehem, R. A., Vértes, P. E., Morgan, S. E., Váša, F., Romero-Garcia, R., Lalonde, F. M., Clasen, L. S., et al. (2020). Transcriptomic and cellular decoding of regional brain vulnerability to neurodevelopmental disorders. Nature Communications, .(.):.

[102] Serdiuk, T., Steudle, A., Mari, S. A., Manioglu, S., Kaback, H. R., Kuhn, A., and Müller, D. J. (2019). Insertion and folding pathways of single membrane proteins guided by translocases and insertases. Science advances, 5(1):eaau6824.

[103] Shafiei, G., Baillet, S., and Misic, B. (2022a). Human electromagnetic and haemodynamic networks systematically converge in unimodal cortex and diverge in transmodal cortex. PLoS biology, 20(8):e3001735.

[104] Shafiei, G., Bazinet, V., Dadar, M., Manera, A. L., Collins, D. L., Dagher, A., Borroni, B., Sanchez-Valle, R., Moreno, F., Laforce, R., et al. (2022b). Network structure and transcriptomic vulnerability shape atrophy in frontotemporal dementia. Brain.

[105] Shafiei, G., Markello, R. D., Makowski, C., Talpalaru, A., Kirschner, M., Devenyi, G. A., Guma, E., Hagmann, P., Cashman, N. R., Lepage, M., et al. (2020a). Spatial patterning of tissue volume loss in schizophrenia reflects brain network architecture. Biological psychiatry, 87(8):727–735.

[106] Shafiei, G., Markello, R. D., Vos de Wael, R., Bernhardt, B. C., Fulcher, B. D., and Misic, B. (2020b). Topographic gradients of intrinsic dynamics across neocortex. elife, 9:e62116.

[107] Sharma, K., Schmitt, S., Bergner, C. G., Tyanova, S., Kannaiyan, N., Manrique-Hoyos, N., Kongi, K., Cantuti, L., Hanisch, U.-K., Philips, M.-A., et al. (2015). Cell type– and brain region–resolved mouse brain proteome. Nature neuroscience, 18(12):1819–1831.

[108] Shen, K., Bezgin, G., Schirner, M., Ritter, P., Everling, S., and McIntosh, A. R. (2019). A macaque connectome for large-scale network simulations in thevirtualbrain. Scientific data, 6(1):123.

[109] Shin, J., French, L., Xu, T., Leonard, G., Perron, M., Pike, G. B., Richer, L., Veillette, S., Pausova, Z., and Paus, T. (2018). Cell-specific gene-expression profiles and cortical thickness in the human brain. Cerebral cortex, 28(9):3267–3277.

[110] Shine, J. M., Breakspear, M., Bell, P. T., Martens, K. A. E., Shine, R., Koyejo, O., Sporns, O., and Poldrack, R. A. (2019). Human cognition involves the dynamic integration of neural activity and neuromodulatory systems. Nat Neurosci, 22(2):289–296.

[111] Stephan, K. E., Kamper, L., Bozkurt, A., Burns, G. A., Young, M. P., and Kötter, R. (2001). Advanced database methodology for the collation of connectivity data on the macaque brain (cocomac). Philosophical Transactions of the Royal Society of London. Series B: Biological Sciences, 356(1412):1159–1186.

[112] Unterholzner, J., Gryglewski, G., Philippe, C., Seiger, R., Pichler, V., Godbersen, G. M., Berroterán-Infante, N., Murgaš, M., Hahn, A., Wadsak, W., et al. (2020). Topologically guided prioritization of candidate gene transcripts coexpressed with the 5-ht1a receptor by combining in vivo pet and allen human brain atlas data. Cerebral cortex, 30(6):3771–3780.

[113] Van Essen, D. C. and Dierker, D. L. (2007). Surface-based and probabilistic atlases of primate cerebral cortex. Neuron, 56(2):209–225.

[114] Vanderhaeghen, P. and Polleux, F. (2023). Developmental mechanisms underlying the evolution of human cortical circuits. Nature Reviews Neuroscience, 24(4):213– 232.

[115] Váša, F. and Mišić, B. (2022). Null models in network neuroscience. Nature Reviews Neuroscience, pages 1–12.

[116] Váša, F., Romero-Garcia, R., Kitzbichler, M. G., Seidlitz, J., Whitaker, K. J., Vaghi, M. M., Kundu, P., Patel, A. X., Fonagy, P., Dolan, R. J., et al. (2020). Conservative and disruptive modes of adolescent change in human brain functional connectivity. Proc Natl Acad Sci USA, 117(6):3248–3253.

[117] Vértes, P., Rittman, T., Whitaker, K., Romero-Garcia, R., Váša, F., Kitzbichler, M., Fonagy, P., Dolan, R., Jones, P., Goodyer, I., et al. (2016). Gene transcription profiles associated with intra-modular and inter-modular hubs in human fmri networks. Philos Trans R Soc Lond B Biol Sci, 371:735–769.

[118] Vogel, C. and Marcotte, E. M. (2012). Insights into the regulation of protein abundance from proteomic and transcriptomic analyses. Nature reviews genetics, 13(4):227–232.

[119] Vos de Wael, R., Benkarim, O., Paquola, C., Lariviere, S., Royer, J., Tavakol, S., Xu, T., Hong, S.-J., Langs, G., Valk, S., et al. (2020). Brainspace: a toolbox for the analysis of macroscale gradients in neuroimaging and connectomics datasets. Communications biology, 3(1):103.

[120] Wagner, H. H. and Dray, S. (2015). Generating spatially constrained null models for irregularly spaced data using moran spectral randomization methods. Methods in Ecology and Evolution, 6(10):1169–1178.

[121] Wagstyl, K., Adler, S., Seidlitz, J., Vandekar, S., Mallard, T. T., Dear, R., DeCasien, A. R., Satterthwaite, T. D., Liu, S., Vértes, P. E., et al. (2024). Transcriptional cartography integrates multiscale biology of the human cortex. Elife, 12:RP86933.

[122] Wagstyl, K., Larocque, S., Cucurull, G., Lepage, C., Cohen, J. P., Bludau, S., Palomero-Gallagher, N., Lewis, L. B., Funck, T., Spitzer, H., et al. (2020). Bigbrain 3d atlas of cortical layers: Cortical and laminar thickness gradients diverge in sensory and motor cortices. PLoS biology, 18(4):e3000678.

[123] Wang, D., Eraslan, B., Wieland, T., Hallström, B., Hopf, T., Zolg, D. P., Zecha, J., Asplund, A., Li, L.-h., Meng, C., et al. (2019). A deep proteome and transcriptome abundance atlas of 29 healthy human tissues. Molecular systems biology, 15(2):e8503.

[124] Warrington, S., Thompson, E., Bastiani, M., Dubois, J., Baxter, L., Slater, R., Jbabdi, S., Mars, R. B., and Sotiropoulos, S. N. (2022). Concurrent mapping of brain ontogeny and phylogeny within a common space: Standardized tractography and applications. Science Advances, 8(42):eabq2022.

[125] Wei, Y., de Lange, S. C., Scholtens, L. H., Watanabe, K., Ardesch, D. J., Jansen, P. R., Savage, J. E., Li, L., Preuss, T. M., Rilling, J. K., et al. (2019). Genetic mapping and evolutionary analysis of human-expanded cognitive networks. Nature communications, 10(1):4839.

[126] Wenthold, R. J., Prybylowski, K., Standley, S., Sans, N., and Petralia, R. S. (2003). Trafficking of nmda receptors. Annual review of pharmacology and toxicology, 43(1):335–358.

[127] Whitaker, K. J., Vértes, P. E., Romero-Garcia, R., Váša, F., Moutoussis, M., Prabhu, G., Weiskopf, N., Callaghan, M. F., Wagstyl, K., Rittman, T., et al. (2016). Adolescence is associated with genomically patterned consolidation of the hubs of the human brain connectome. Proceedings of the National Academy of Sciences, 113(32):9105–9110.

[128] Xu, T., Nenning, K. H., Schwartz, E., Hong, S. J., Vogelstein, J. T., Goulas, A., Fair, D. A., Schroeder, C. E., Margulies, D. S., Smallwood, J., Milham, M. P., and Langs, G. (2020). Cross-species functional alignment reveals evolutionary hierarchy within the connectome. NeuroImage, 223. PMID: 32916286 publisher: Academic Press Inc.

[129] Yang, S.-H., Cheng, P.-H., Banta, H., Piotrowska-Nitsche, K., Yang, J.-J., Cheng, E. C., Snyder, B., Larkin, K., Liu, J., Orkin, J., et al. (2008). Towards a transgenic model of huntington’s disease in a non-human primate. Nature, 453(7197):921–924.

[130] Yudowski, G. A., Puthenveedu, M. A., and von Zastrow, M. (2006). Distinct modes of regulated receptor insertion to the somatodendritic plasma membrane. Nature neuroscience, 9(5):622–627.

[131] Zachlod, D., Bludau, S., Cichon, S., Palomero-Gallagher, N., and Amunts, K. (2022). Combined analysis of cytoarchitectonic, molecular and transcriptomic patterns reveal differences in brain organization across human functional brain systems. NeuroImage, 257:119286.

[132] Zarkali, A., Luppi, A. I., Stamatakis, E. A., Reeves, S., McColgan, P., Leyland, L.-A., Lees, A. J., and Weil, R. S. (2022). Changes in dynamic transitions between integrated and segregated states underlie visual hallucinations in parkinson’s disease. Communications Biology, 5(1):928.

[133] Zarkali, A., McColgan, P., Ryten, M., Reynolds, R., Leyland, L.-A., Lees, A. J., Rees, G., and Weil, R. S. (2020). Differences in network controllability and regional gene expression underlie hallucinations in parkinson’s disease. Brain, 143(11):3435–3448.

[134] Zhang, B., Wang, J., Wang, X., Zhu, J., Liu, Q., Shi, Z., Chambers, M. C., Zimmerman, L. J., Shaddox, K. F., Kim, S., et al. (2014a). Proteogenomic characterization of human colon and rectal cancer. Nature, 513(7518):382– 387.

[135] Zhang, Y., Chen, K., Sloan, S. A., Bennett, M. L., Scholze, A. R., O’Keeffe, S., Phatnani, H. P., Guarnieri, P., Caneda, C., Ruderisch, N., et al. (2014b). An rna-sequencing transcriptome and splicing database of glia, neurons, and vascular cells of the cerebral cortex. Journal of Neuroscience, 34(36):11929–11947.

[136] Zhao, L., Mühleisen, T. W., Pelzer, D. I., Burger, B., Beins, E. C., Forstner, A. J., Herms, S., Hoffmann, P., Amunts, K., Palomero-Gallagher, N., et al. (2023). Relationships between neurotransmitter receptor densities and expression levels of their corresponding genes in the human hippocampus. Neuroimage, 273:120095.

