## Supplementary Material for "Benchmarking macaque brain gene expression for horizontal and vertical translation" (from another version of this preprint: version 1, 2024-08-19)

**Supplementary Figures**

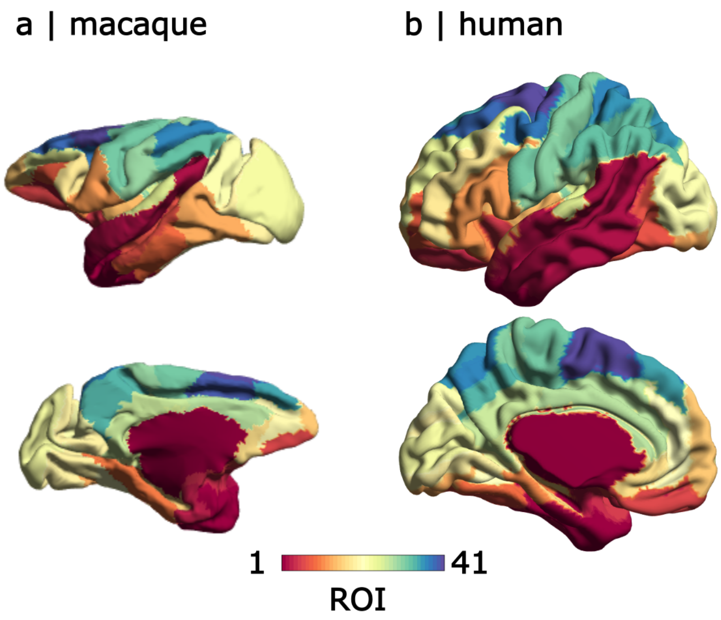


Figure S1. **Regional Mapping parcellation for different species |** (**a**) The macaque cortical atlas of Kötter and Wanke [66]. (**b**) Human translation of the same atlas by [10].

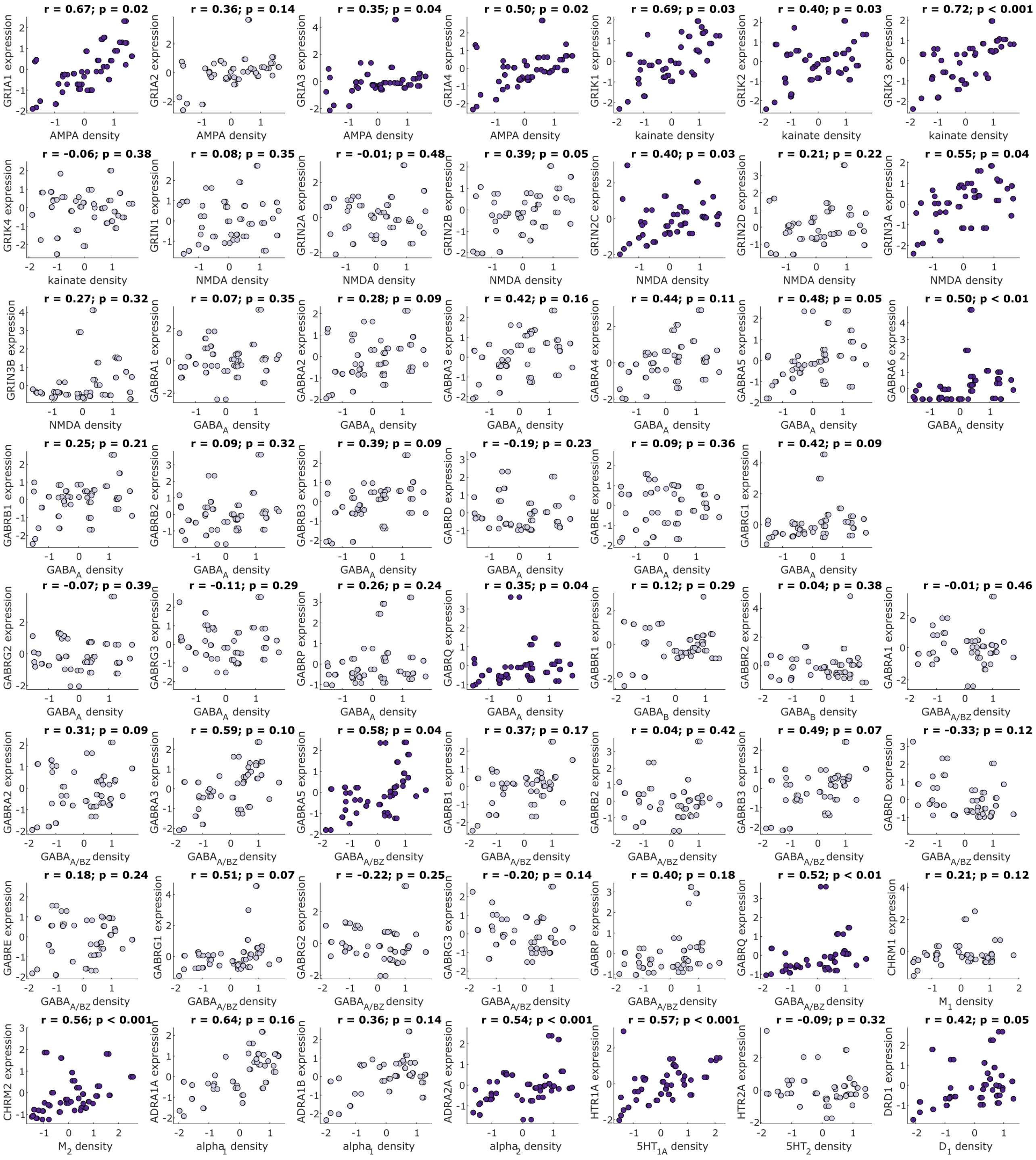


Figure S2. **Autoradiography-derived receptor densities versus gene expression in the macaque, for all pairs |** The density of only three neurotransmitter receptors correlates significantly with the expression of their corresponding gene, across macaque cortex. Indigo scatter plots indicate significant human-macaque correspondence (Spearman's *r*, $p < 0.05$ against a null distribution with preserved spatial autocorrelation). Values are z-scored.

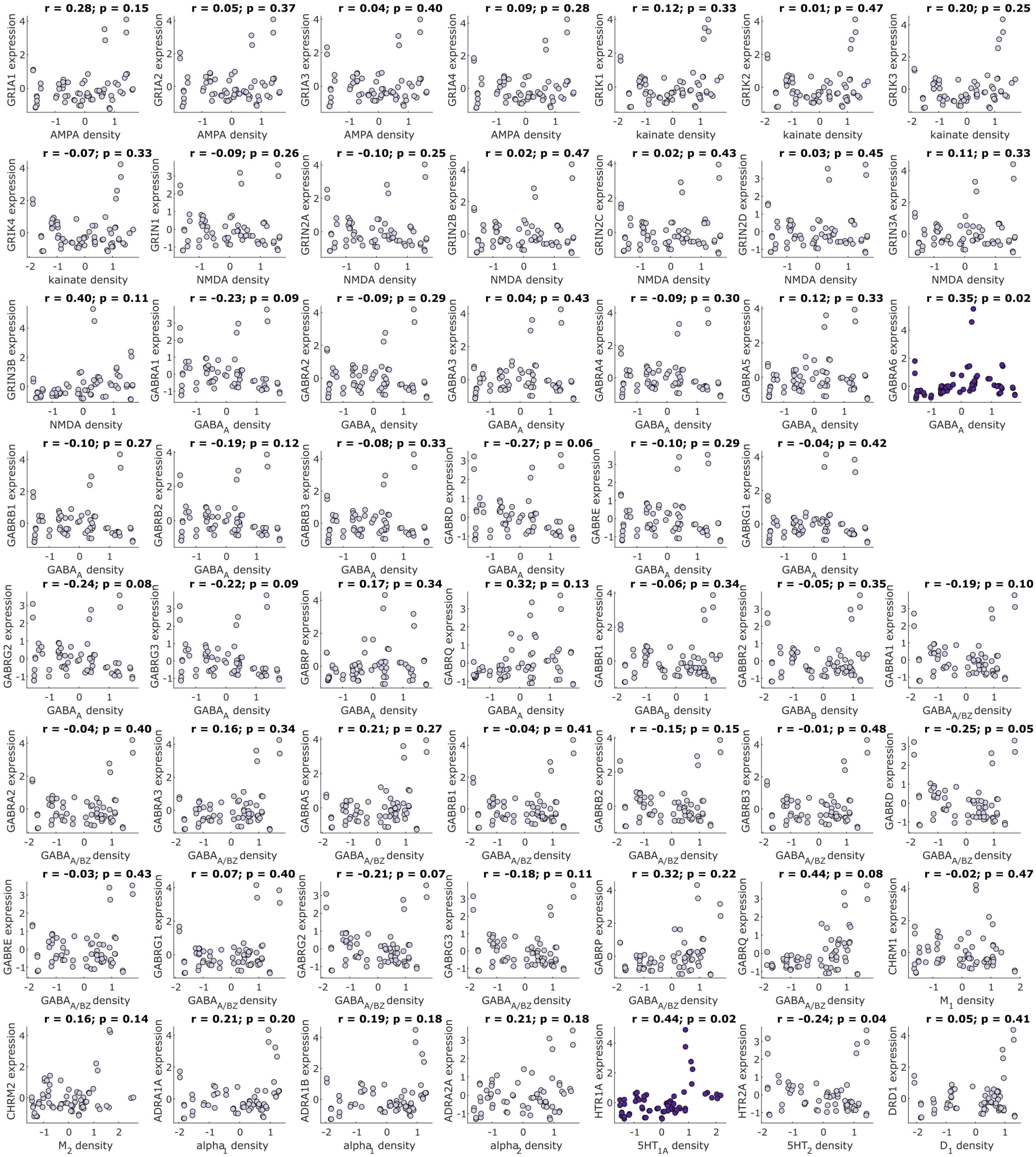


Figure S3. **Autoradiography-derived receptor densities versus mean gene expression of each region's neighbours, weighted by efferent connection strength |** Indigo scatter plots indicate significant human-macaque correspondence (Spearman's $r$, $p < 0.05$ against a null distribution with preserved spatial autocorrelation). Values are z-scored.

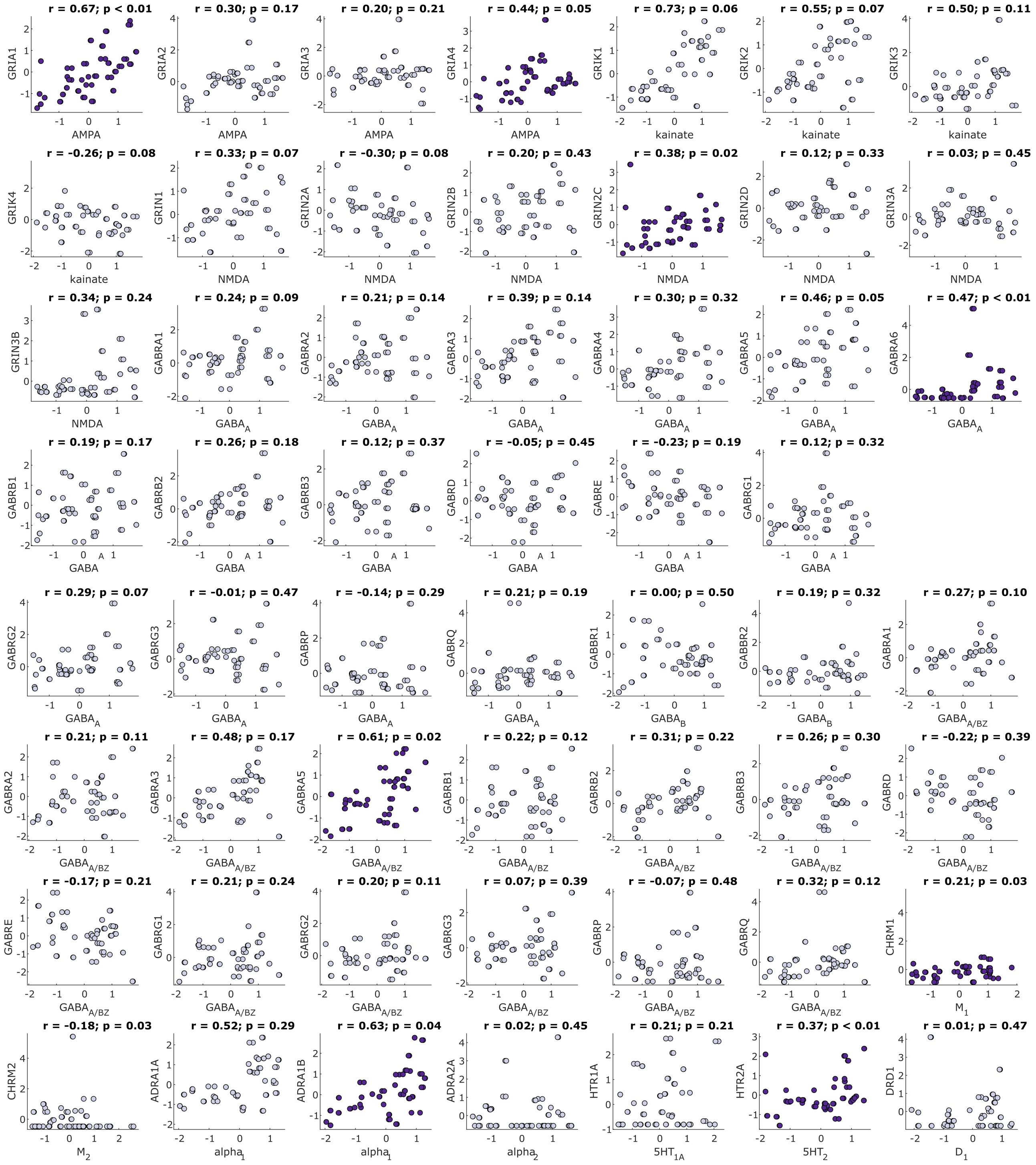


Figure S4. **Autoradiography-derived receptor densities versus Layer 1 gene expression in the macaque, for all pairs |** The density of only three neurotransmitter receptors correlates significantly with the expression of their corresponding gene, across macaque cortex. Indigo scatter plots indicate significant human-macaque correspondence (Spearman's $r$, $p < 0.05$ against a null distribution with preserved spatial autocorrelation). Values are z-scored.

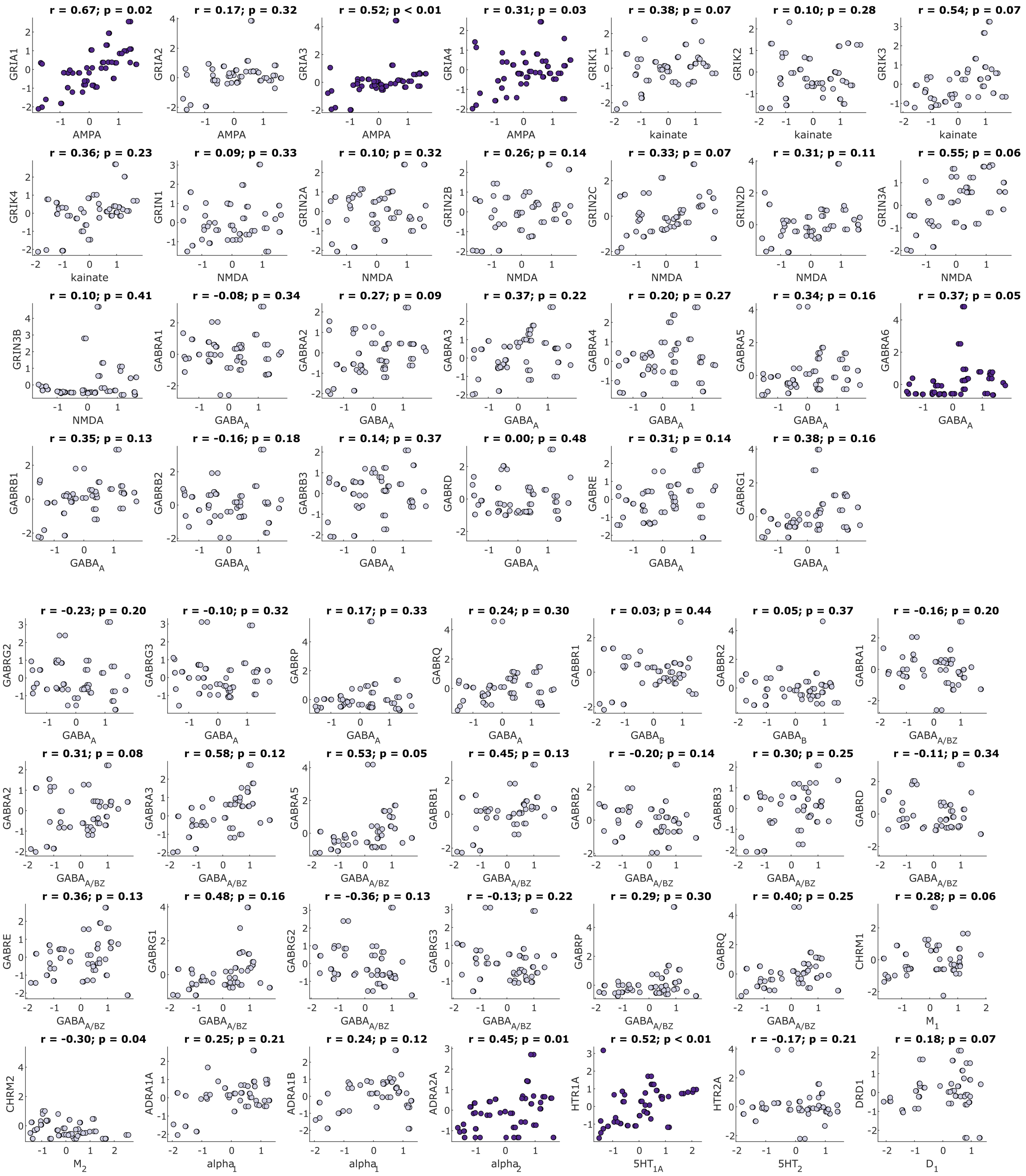


Figure S5. **Autoradiography-derived receptor densities versus Layer 2 gene expression in the macaque, for all pairs |** The density of only three neurotransmitter receptors correlates significantly with the expression of their corresponding gene, across macaque cortex. Indigo scatter plots indicate significant human-macaque correspondence (Spearman's *r*, $p < 0.05$ against a null distribution with preserved spatial autocorrelation). Values are z-scored.

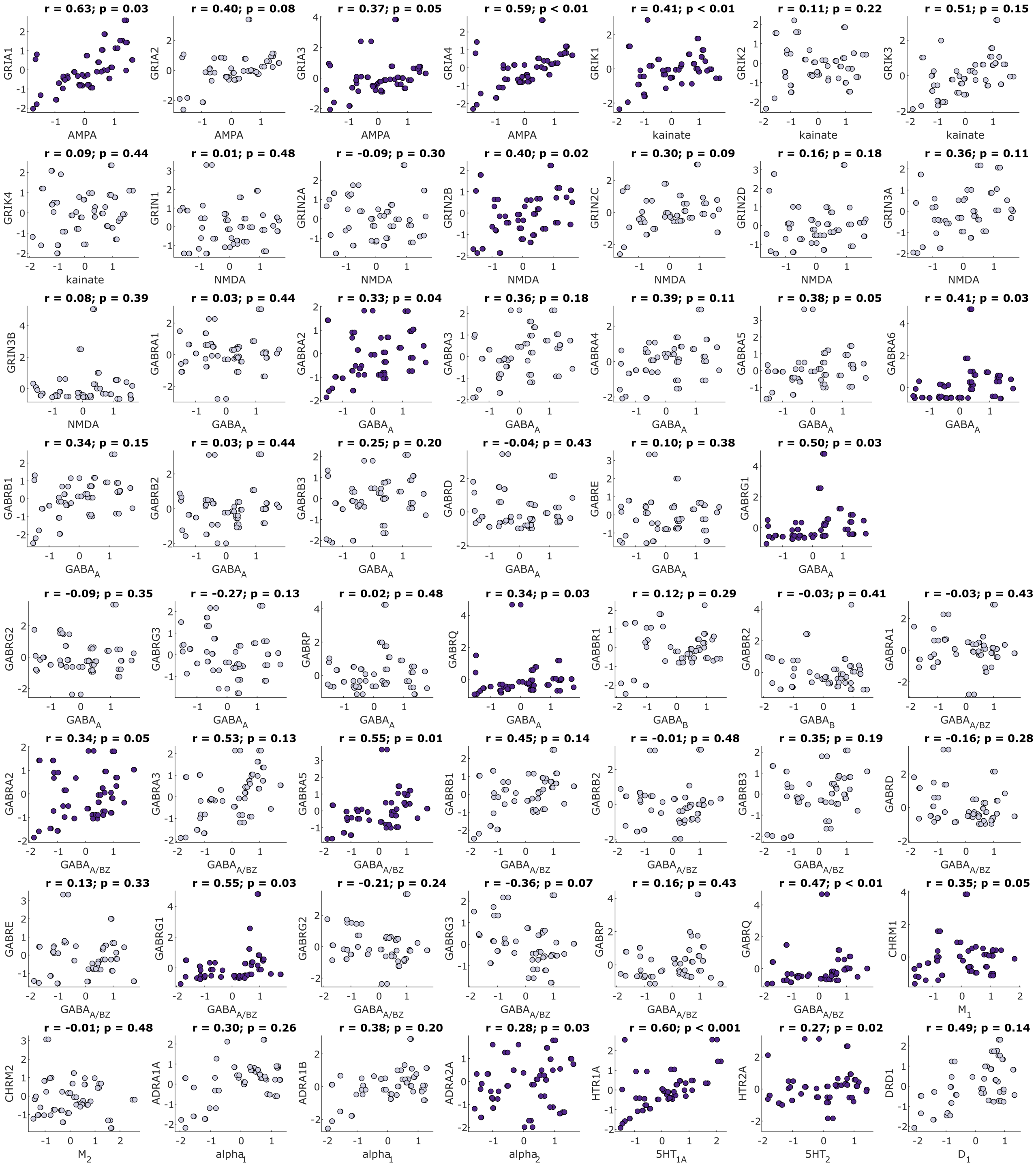


Figure S6. **Autoradiography-derived receptor densities versus Layer 3 gene expression in the macaque, for all pairs |** The density of only three neurotransmitter receptors correlates significantly with the expression of their corresponding gene, across macaque cortex. Indigo scatter plots indicate significant human-macaque correspondence (Spearman's $r$, $p < 0.05$ against a null distribution with preserved spatial autocorrelation). Values are z-scored.

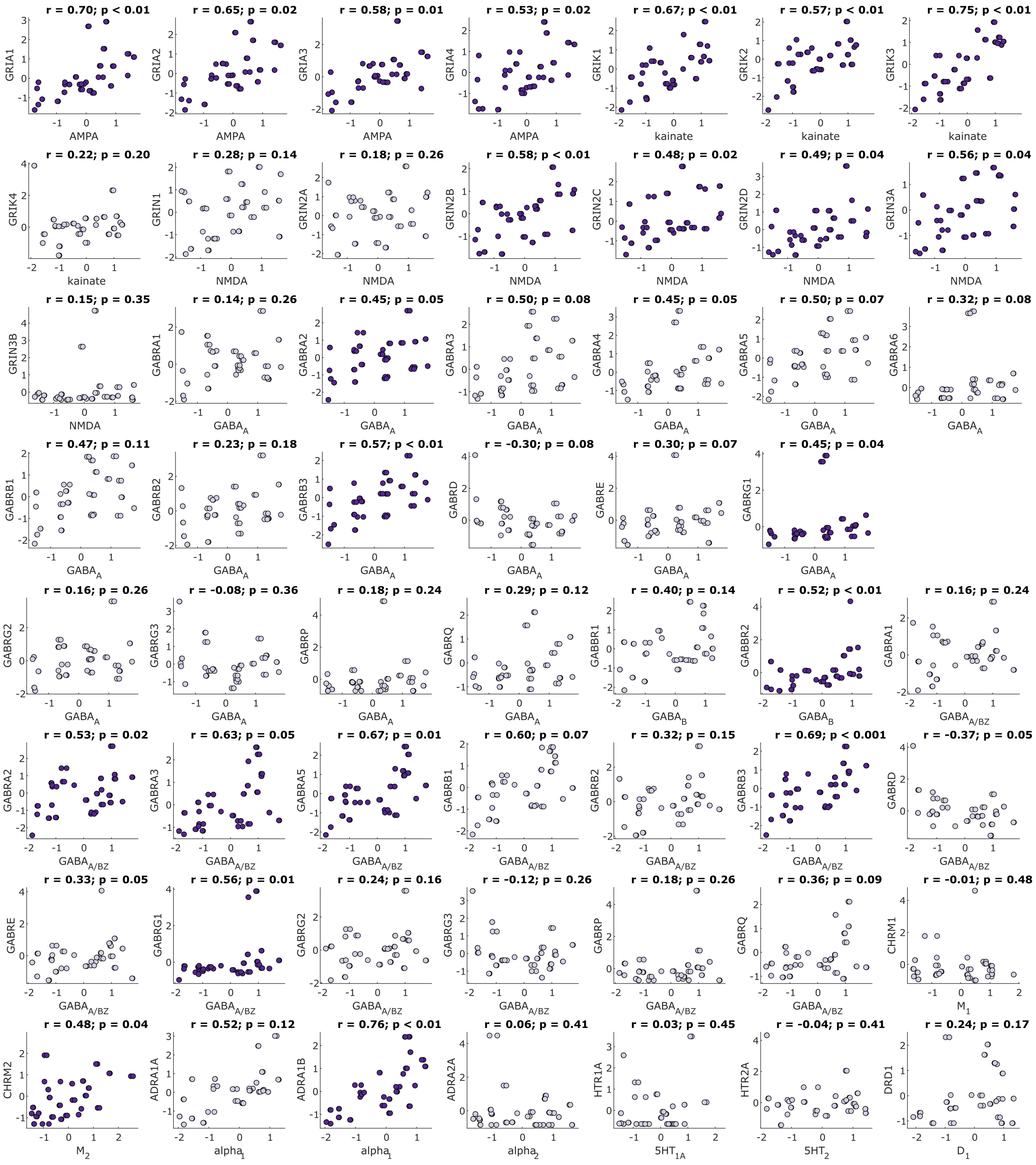


Figure S7. **Autoradiography-derived receptor densities versus Layer 4 gene expression in the macaque, for all pairs |** The density of only three neurotransmitter receptors correlates significantly with the expression of their corresponding gene, across macaque cortex. Indigo scatter plots indicate significant human-macaque correspondence (Spearman's $r$, $p < 0.05$ against a null distribution with preserved spatial autocorrelation). Values are z-scored.

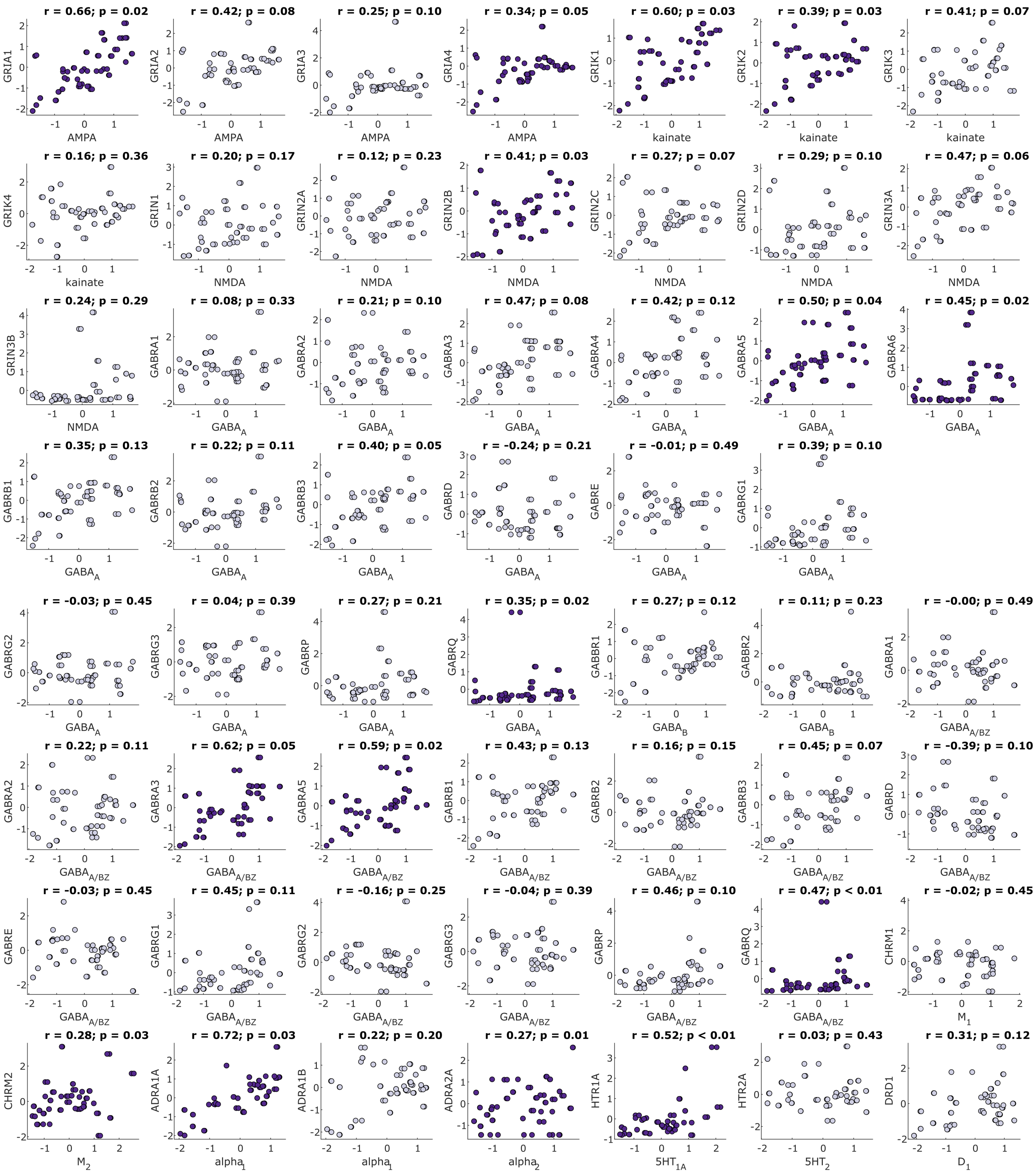


Figure S8. **Autoradiography-derived receptor densities versus Layer 5 gene expression in the macaque, for all pairs |** The density of only three neurotransmitter receptors correlates significantly with the expression of their corresponding gene, across macaque cortex. Indigo scatter plots indicate significant human-macaque correspondence (Spearman's $r$, $p < 0.05$ against a null distribution with preserved spatial autocorrelation). Values are z-scored.

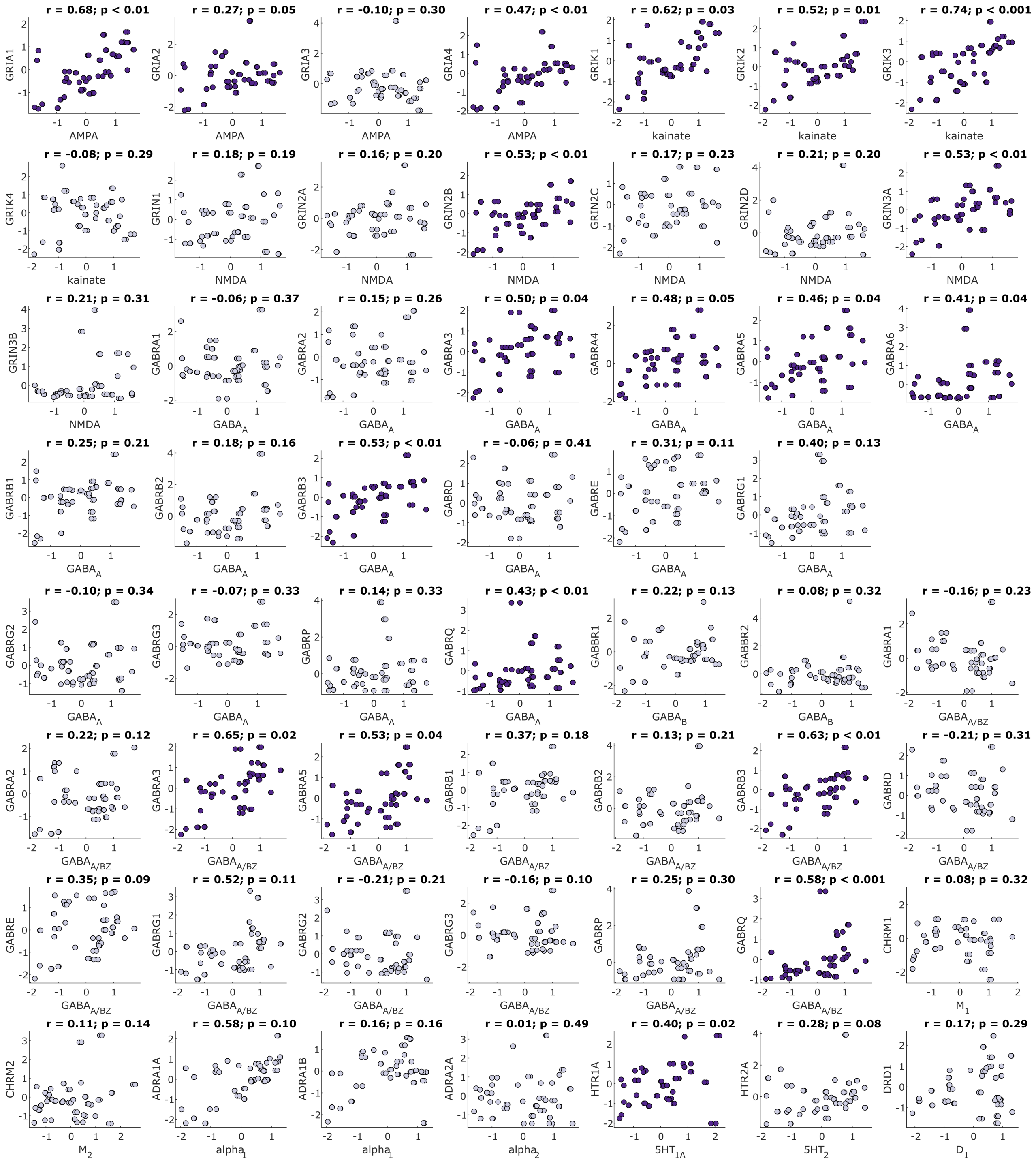

Figure S9. **Autoradiography-derived receptor densities versus Layer 6 gene expression in the macaque, for all pairs |** The density of only three neurotransmitter receptors correlates significantly with the expression of their corresponding gene, across macaque cortex. Indigo scatter plots indicate significant human-macaque correspondence (Spearman's $r$, $p < 0.05$ against a null distribution with preserved spatial autocorrelation). Values are z-scored.

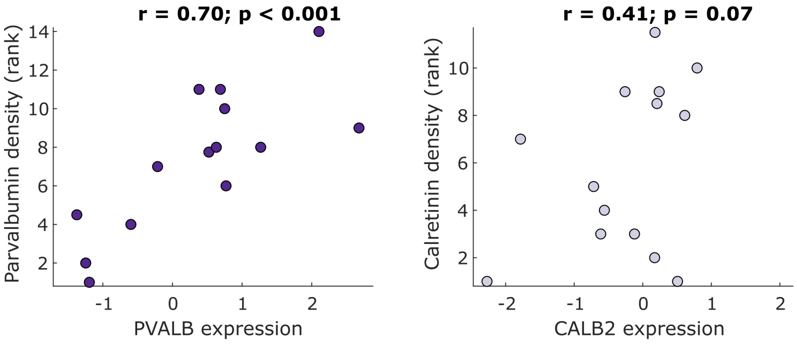


Figure S10. **Spatial correspondence of macaque parvalbumin- and calretinin-immunoreactive neurons with gene expression, for a subset of visual, auditory, and somatosensory areas |** Left: regional rank of parvalbumin-immunoreactive neurons versus regional *PVALB* gene expression. Right: regional rank of calretinin-immunoreactive neurons versus regional *CALB2* gene expression. Indigo scatter plots indicate significant spatial correspondence (Spearman's $r$, $p < 0.05$ against a null distribution with preserved spatial autocorrelation). Values are z-scored.

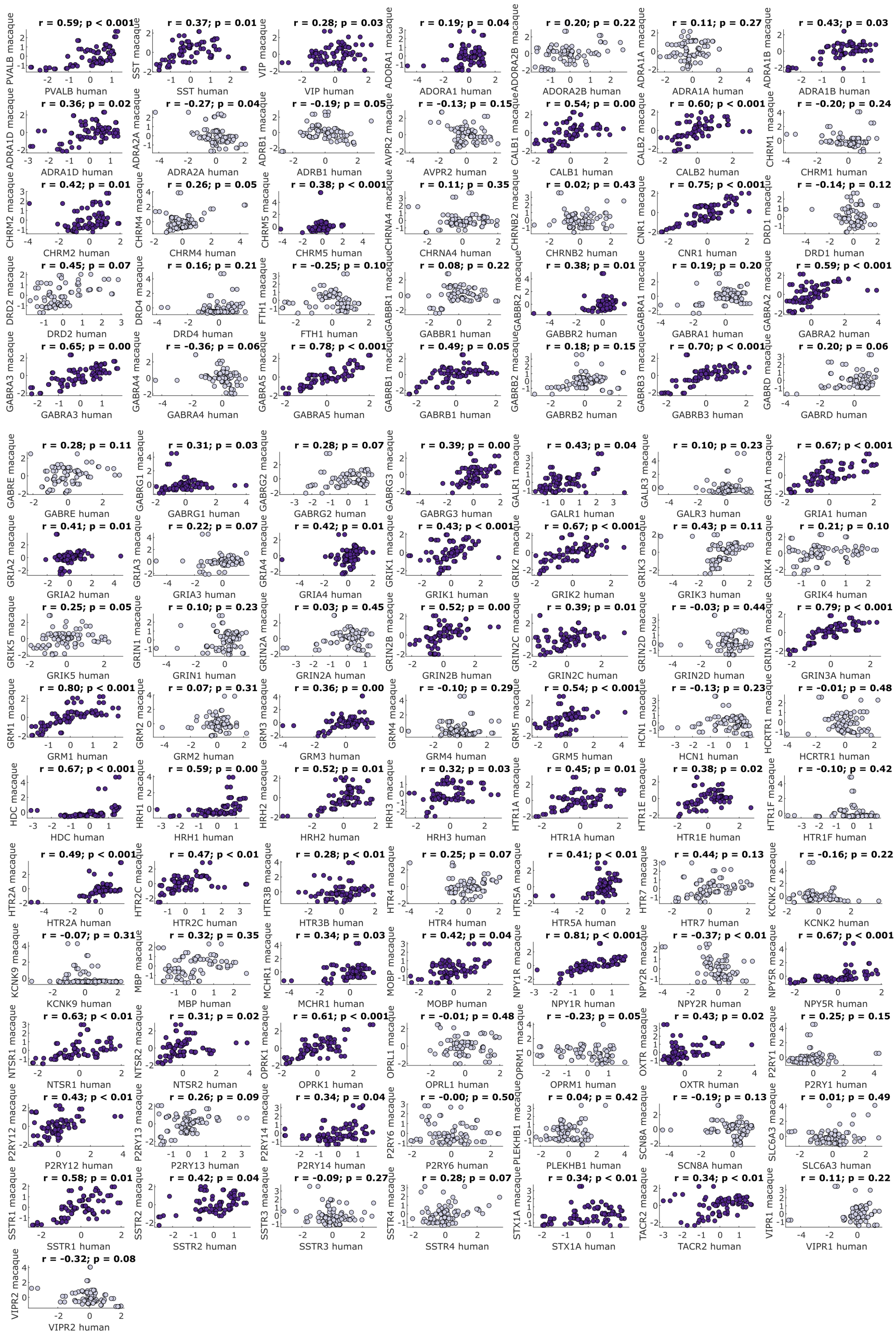


Figure S11. **Spatial correspondence between human gene expression from microarray data and macaque gene expression from stereo-seq, for all brain-related genes considered.** Indigo scatter plots indicate significant ($p < 0.05$) human-macaque correspondence. Gene expression is z-scored.

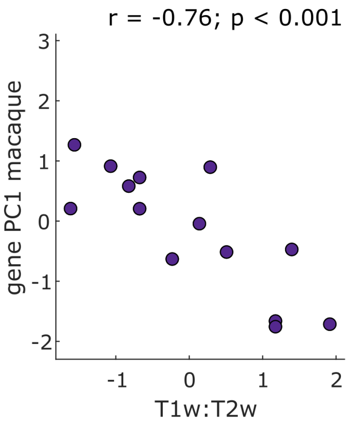


Figure S12. **Replication of spatial correspondence between macaque gene PC1 and intracortical myelination |** Abscissa: intracortical myelination (T1w:T2w ratio) from [16]. Ordinate: macaque gene PC1. Values are z-scored. Significance of correlation is assessed against a null distribution with preserved spatial autocorrelation.

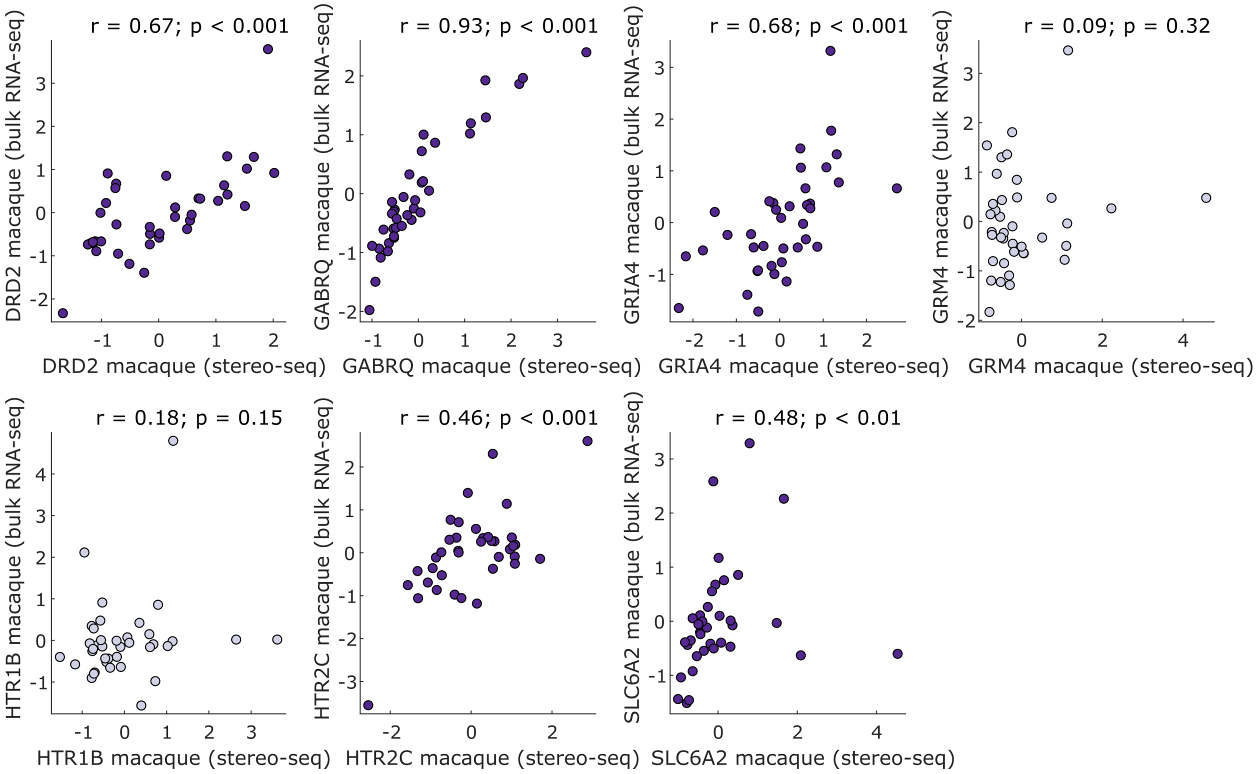


Figure S13. **Spatial correspondence between macaque gene expression datasets |** Abscissa: macaque gene expression from stereo-seq [22]. Ordinate: macaque gene expression from RNA-seq [12]. Indigo scatter plots indicate significant ($p < 0.05$) correspondence. Gene expression is z-scored.

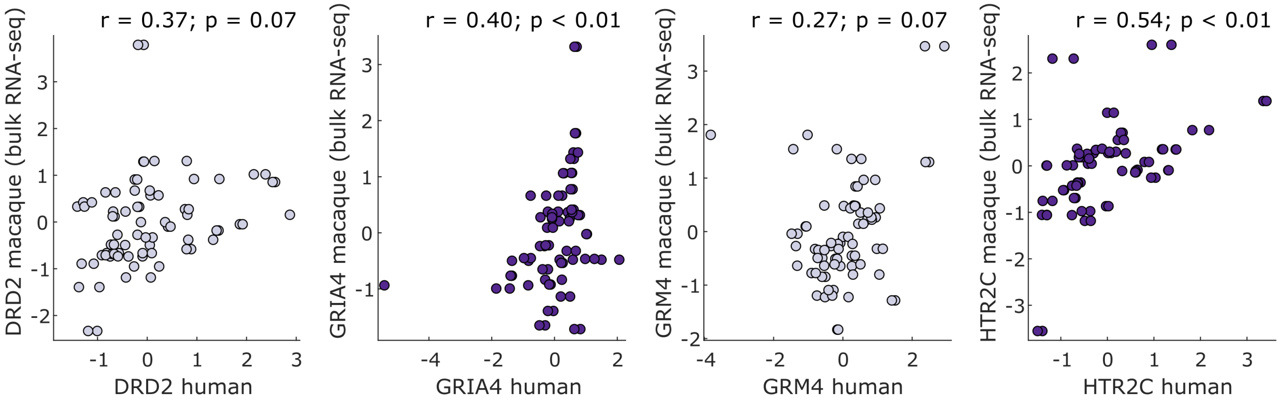


Figure S14. **Spatial correspondence between human gene expression from AHBA microarray and macaque gene expression from RNA-seq |** Abscissa: human gene expression from microarray data [54]. Ordinate: macaque gene expression from RNA-seq [12]. Indigo scatter plots indicate significant ($p < 0.05$) human-macaque correspondence. Gene expression is z-scored.

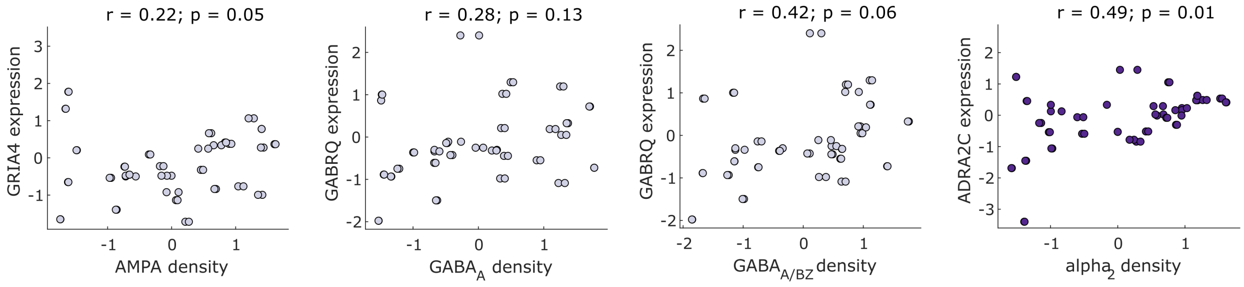


Figure S15. **Spatial correspondence between macaque receptor density and macaque gene expression from RNA-seq |** Abscissa: macaque receptor density from autoradiography [39]. Ordinate: macaque gene expression from RNA-seq [12]. Indigo scatter plots indicate significant ($p < 0.05$) gene-receptor correspondence. Values are z-scored.

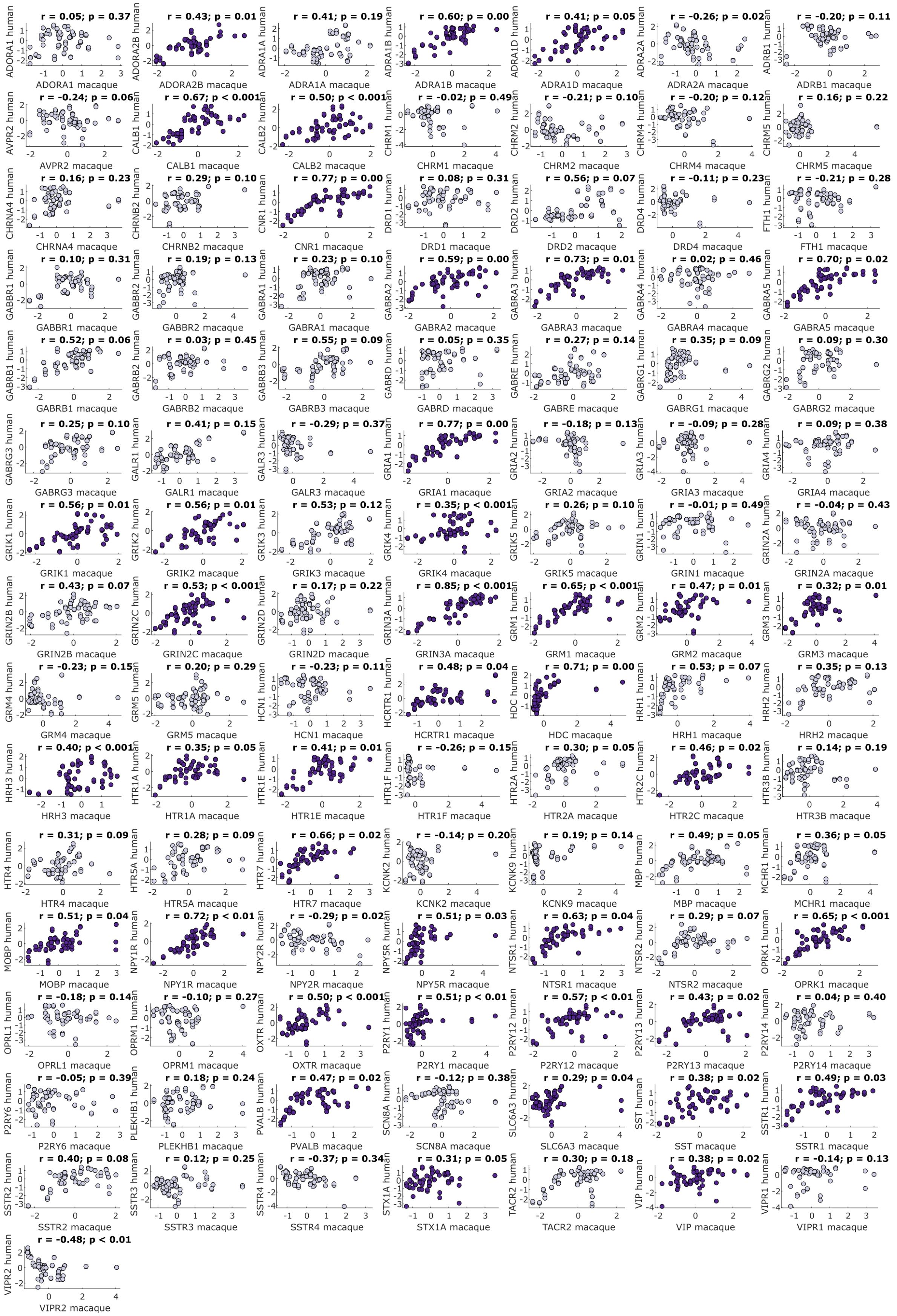


Figure S16. **Spatial correspondence between human gene expression from RNA-seq and macaque gene expression from stereo-seq |** Indigo scatter plots indicate significant ($p < 0.05$) human-macaque correspondence. Gene expression is z-scored.

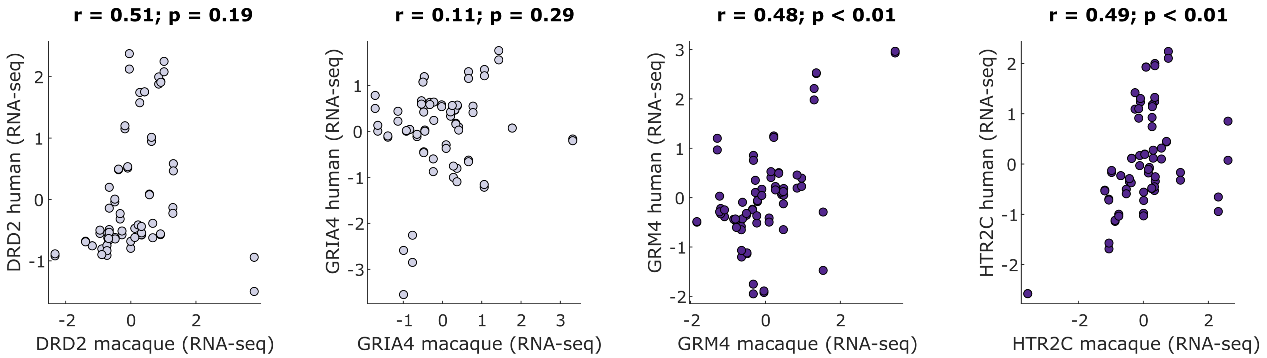


Figure S17. **Spatial correspondence between human gene expression from RNA-seq and macaque gene expression from RNA-seq |** Abscissa: human gene expression from RNA-seq data [54]. Ordinate: macaque gene expression from RNA-seq [12]. Indigo scatter plots indicate significant ($p < 0.05$) human-macaque correspondence. Gene expression is z-scored.

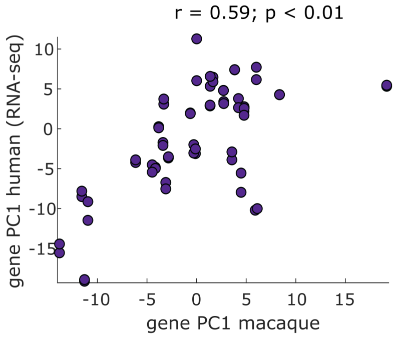


Figure S18. **Spatial correspondence of macaque gene PC1 from stereo-seq with human gene PC1 from RNA-seq |** Abscissa: macaque gene PC1 from stereo-seq. Ordinate: human gene PC1 from RNA-seq. Values are z-scored. Significance of correlation is assessed against a null distribution with preserved spatial autocorrelation.

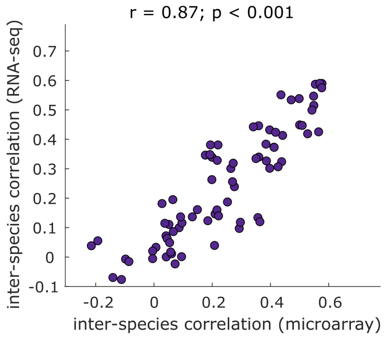


Figure S19. **The pattern of regional inter-species correlation of gene expression is recapitulated when using human RNA-seq data |** Abscissa: regional inter-species correlation of gene expression, using macaque stereo-seq and human microarray data. Ordinate: regional inter-species correlation of gene expression, using macaque stereo-seq and human RNA-seq data. Significance of correlation is assessed against a null distribution with preserved spatial autocorrelation.

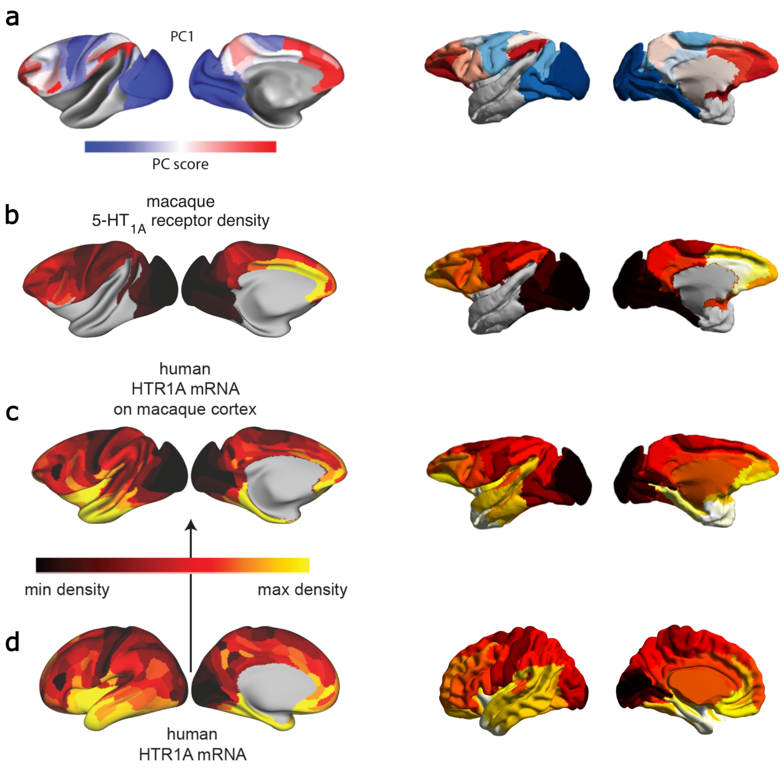


Figure S20. **Mapping macaque receptor autoradiography and human gene expression data to the Regional Mapping parcellation |** (**a**) PC1 of macaque receptor density from autoradiography, in the original parcellation from [39] (left) and mapped onto the macaque Regional Mapping atlas of [66] (right). (**b**) Macaque *5HT*$_{1A}$ receptor density, in the original parcellation from [39] (left) and mapped onto the macaque Regional Mapping atlas of [66] (right). (**c**) Human *HTR1A* mRNA mapped onto the macaque parcellation by [39] (left) and mapped onto the macaque Regional Mapping atlas of [66] (right). (**d**) Human *HTR1A* mRNA, mapped on the HCP-MMP human cortical parcellation of [44] by [39] (left) and mapped onto the human version of the Regional Mapping atlas made by [10] (right). Adapted from Figures 2 and 7 of [39], published under CC-BY license.
